# Extracellular matrix regulates lineage plasticity in prostate cancer through YAP/TEAD

**DOI:** 10.64898/2025.12.30.697072

**Authors:** Teng Han, Zhen Sun, Matthew Lange, Y Zoe Cho, Patrick Mcgillivray, Maren Büttner, Nathaniel R Kastan, Subhiksha Nandakumar, Huiyong Zhao, Eralda Salataj, Sanyukta Oak, Linda Fong, Wenfei Kang, Ning Fan, Francesca Mazzoni, Radhika A Patel, Jimmy Zhao, Nazifa Salsabeel, Harmanpreet Kaur, Ninghui Mao, Qing Chang, Eric Rosiek, Eric Chan, Murray Tipping, Nikolaus Schultz, Pierre-Jacques Hamard, Elisa DeStanchina, Zhenghao Chen, Michael C Haffner, Dana Pe’er, Richard Koche, A James Hudspeth, Charles L Sawyers

**Author notes:** Corresponding author: Charles L Sawyers. SystImmune Inc, Redmond, WA 98052, USA. Deceased (A James Hudspeth).

## Abstract

Treatment-related neuroendocrine prostate cancer (NEPC) is an increasingly frequent mechanism of resistance to androgen receptor pathway inhibitor (ARPI) therapy in prostate adenocarcinoma (PRAD). This lineage transition is dependent on upregulation of the NE-specifying transcription factor ASCL1, typically in a genetic background of *RB1* and *TP53* loss. Here we identify extracellular matrix-integrin-YAP1/TEAD signaling as a critical brake on NEPC lineage transition. Deletion of *Itgb1*, the shared B1 subunit required for collagen and laminin-mediated integrin activation, is sufficient to induce ASCL1 and NE lineage gene expression, by activating LATS1/2 kinases with subsequent inactivation of YAP1/TEAD signaling. Conversely, restoration of YAP1/TEAD signaling by pharmacological LATS1/2 inhibition, or by expression of constitutively active YAP1/TAZ mutants, prevents or reverts NEPC lineage transition. NOTCH and AR cooperate with YAP/TEAD to repress ASCL1, such that combined inhibition leads to complete reprogramming of PRAD into NEPC *in vitro*, providing a dynamic platform to dissect the molecular events responsible for lineage transition over time. We find that lineage transition is accompanied by a redistribution of FOXA1 and TEAD cistromes from PRAD to NEPC-specific enhancers and requires the pioneering activity of FOXA1. Thus, extracellular matrix/integrin signaling in the PRAD tumor microenvironment restrains NE lineage plasticity, highlighting a potential path for pharmacological inhibitors in modulating treatment-induced lineage change.

## Introduction

Lineage plasticity - the ability of cancer cells to adopt alternative identities in response to environmental or therapeutic pressures - is increasingly recognized as a major mechanism of resistance to targeted therapies across multiple cancer types, including prostate cancer (*1*, *2*). The mainstay of prostate cancer treatment is androgen deprivation therapy (ADT), to which most patients initially respond. However, resistance inevitably develops, leading to the emergence of castration-resistant prostate cancer (CRPC) (*3*). To counteract reactivated androgen receptor (AR) signaling in CRPC, next-generation AR pathway inhibitors (ARPIs) such as enzalutamide and abiraterone have become standard of care in CRPC and are now used in earlier treatment settings (*4*, *5*). Despite improved overall survival, an unintended consequence of ARPI therapy is selective pressure that results in an increase in treatment-related neuroendocrine prostate cancer (NEPC) from prostate adenocarcinoma (PRAD), a lineage transition resembling EGFR-mutant lung adenocarcinomas that transition to small cell lung cancer (SCLC) following treatment with EGFR inhibitors (*6*, *7*). Once this lineage switch occurs, NEPC is associated with poor clinical outcomes and is largely unresponsive to existing treatments (*8*), underscoring the need to understand the molecular events that initiate lineage plasticity, with the goal of preventing or delaying the transition.

Genomic loss of the *RB1* and *TP53* tumor suppressors is highly enriched in human NEPC (and in EGFR-mutant lung cancers that transition to SCLC) and is required for NEPC lineage transition in mouse models (*9–12*). However, the presence of these mutations alone is not sufficient to initiate lineage reprogramming. For example, *Rb1*/*Trp53* deficient mouse prostate organoids retain a luminal/basal PRAD lineage when propagated *in vitro* but undergo lineage transition after *in vivo* transplantation, suggesting a critical role for the tumor microenvironment (TME) (*9*, *13*). Additionally, genomically annotated cohorts of CRPC and EGFR-mutant lung cancer reveal that not all adenocarcinomas with *RB1* and *TP53* loss undergo lineage transition in response to therapy (*12*, *14*). The fact that treatment-induced lineage transitions can be reversible further suggests that epigenetic programs likely play a critical role (*15–17*). Current evidence supports a model in which *RB1* and *TP53* loss establishes a chromatin state permissive for lineage switching, but additional tumor cell-extrinsic signals from the TME are needed to initiate and sustain lineage plasticity. Here we explore the nature of these TME cues and how they integrate with tumor-intrinsic programs.

## Results

As a first step, we confirmed prior work (*9*, *13*) demonstrating that genetically engineered murine organoids with *Rb1* and *Trp53* loss coupled with *cMyc* overexpression (RPM) undergo a PRAD to NEPC transition following orthotopic transplantation into mice but fail to do so in organoid culture, as indicated by the absence of staining for the NE lineage marker *Ascl1* (**Fig. S1A-B**). We extended this phenotype to two additional *Rb1*/*Trp53*-deficient genotypes: *Rb1^-/-^; Trp53^-/-^; Pten^-/-^* (TKO) and *Rb1^-/-^; Trp53^-/-^; Pten^-/-^; cMyc^+^*(TKOM) (**Fig. S1A-B**). These findings document that conventional organoid culture systems fail to fully replicate the *in vivo* TME necessary for adenocarcinoma-to-NE transition.

### Relative depletion of stroma and extracellular matrix in NEPC

To search for TME factors responsible for this lineage transition, we analyzed histologic sections from wild-type normal prostate tissue, as well as PRAD and NEPC regions from the PtRP (*Pten^-/-^*, *Rb1^-/-^*, *Trp53^-/-^*, *Pb-Cre*) genetically engineered mouse model (GEMM) (*17*). Strikingly, we observed that v*imentin*-positive fibroblasts, which are sparse in normal prostate tissue but expanded in PRAD regions, are relatively depleted in NEPC regions. Similarly, extracellular matrix (ECM) proteins, such as fibronectin and collagen—primarily produced by fibroblasts—are scarce in normal prostate tissue, abundant in PRAD and depleted in NEPC (**Fig. 1A**). Tumor-associated fibroblasts and collagen were also abundant in the TKO and RPM orthotopic transplantation models at the PRAD stage (2 weeks post-transplantation) but depleted following the transition to NEPC (16 weeks for TKO and 7 weeks for RPM) (**Fig. S1C**).

**Fig. 1.**
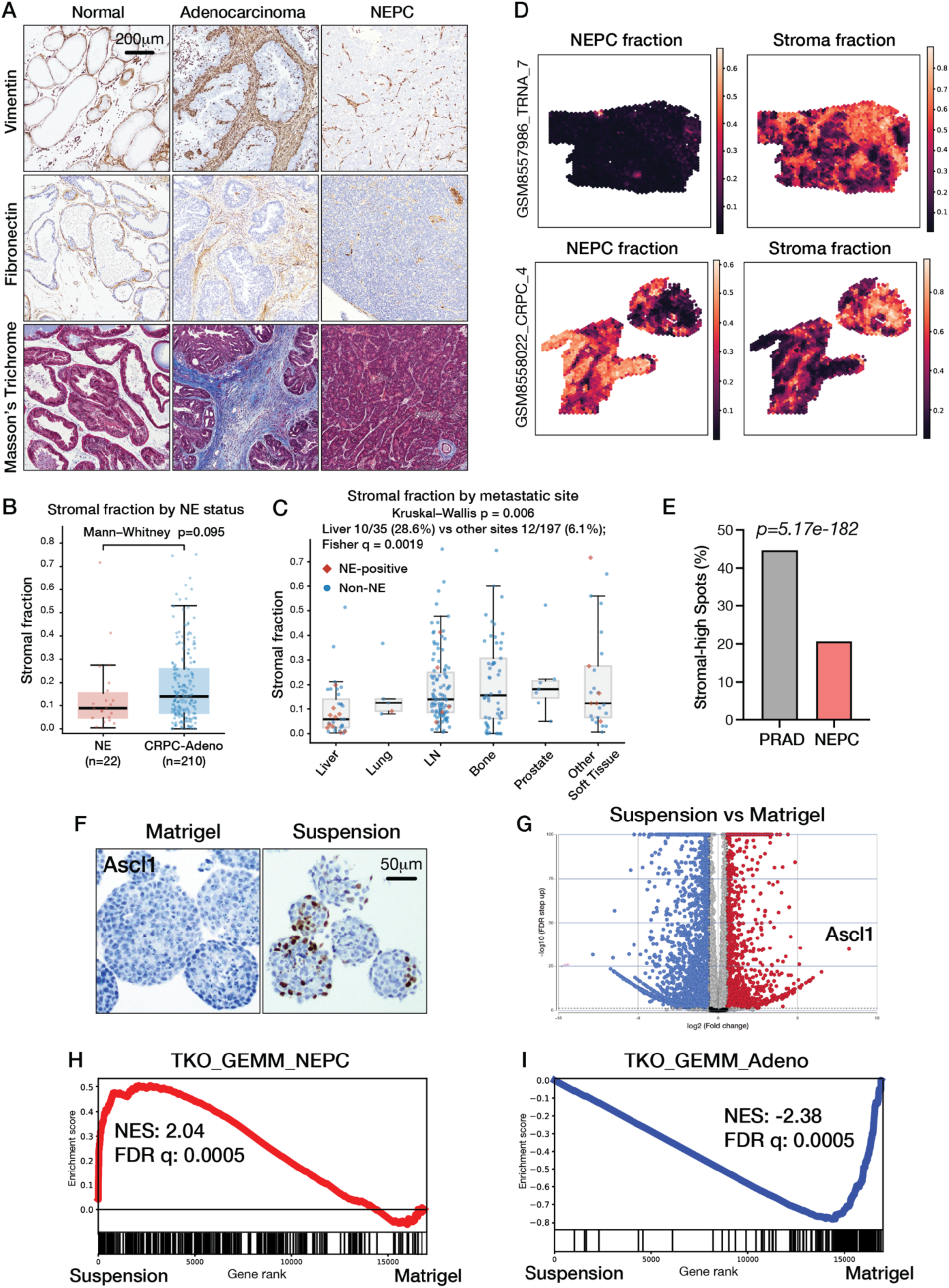
Removal of ECM activates Ascl1 expression. (A) IHC of Vimentin and Fibronectin, along with Masson’s trichrome staining of normal prostate tissue, adenocarcinoma regions, and neuroendocrine regions from TKO tumors. (B) Quantification of stromal fraction in NE-positive (n=73) versus CRPC-adeno (n=210) samples from SU2C/PCF metastatic prostate cancer cohort. Box plots show median and interquartile range; whiskers extend to 1.5X IQR. Mann-Whitney U test, p = 0.095. (C) Quantification of stromal fraction per sample stratified by metastatic site. Individual samples are colored by pathologist-annotated neuroendocrine status (red diamonds, NE-positive; blue circles, non-NE). NE-positive samples are concentrated at stroma-low sites (liver, lung, lymph node), with no NE-positive samples identified among bone (n=73) or prostate (n=7) sites. Fisher q test compares NE prevalence in liver versus all other sites of disease. The Kruskal–Wallis p is a global test showing that stromal fraction differs across sites of metastatic disease (D) Deconvolved NEPC theta estimates and corresponding stromal theta estimates across two specimens. Spatial maps highlight the distribution of NEPC and fibroblast-like content across Visium spots. (E) Proportion of spots classified as both NEPC stromal-high (771/3,719 = 20.7%) and AR/PRAD stromal-high (33,015/73,827 = 44.7%) (χ² p = 5.17 × 10⁻¹⁸²). (F) ASCL1 IHC of TKOM organoids cultured in Matrigel versus suspension conditions (30 days in suspension). (G) Volcano plot showing differentially expressed genes from RNA-seq analysis comparing TKOM organoids cultured in suspension for 30 days versus Matrigel. (H) Gene set enrichment analysis (GSEA) using NEPC signature derived from TKO GEMM of RNA-seq data from TKOM organoids in suspension versus Matrigel culture. The NEPC gene set is listed in table S2. (I) Gene set enrichment analysis (GSEA) using prostate adenocarcinoma signature derived from TKO GEMM of RNA-seq data from TKOM organoids in suspension versus Matrigel culture. The prostate adenocarcinoma (PRAD) gene set is listed in table S2.

To explore whether relative depletion of fibroblasts is also observed in human NEPC, we performed BayesPrism deconvolution on bulk RNA-seq data from the SU2C/PCF metastatic CRPC cohort (n=266) to estimate the fractional contribution of PRAD, NEPC and stroma (**Fig S2A,** see methods). Our estimate of NE status was concordant with published clinical annotations, confirming the utility of this approach (see **Fig S2A** legend). As with the mouse data, NE-positive samples had a reduced median stromal fraction relative to PRAD but this did not reach statistical significance (median theta 0.089 vs 0.142, p = 0.095) (**Fig 1B**). When stratified by metastatic site, NEPC showed a clear predilection for stroma-poor (liver, lymph node, lung) versus stroma-rich (bone) sites, consistent with known clinical metastasis patterns (**Fig 1C**). To examine the PRAD/NEPC/stroma question from a spatial perspective, we used BayesPrism deconvolution to estimate the fractional contribution of each cell type within each Visium spot of a recently published prostate cancer cohort profiled by spatial transcriptomics (*18–20*) (**Fig S2A**, see methods). Among PRAD-high spots, 44.7% had high stromal content (33,015/73,827) compared to 20.7% of NEPC-high spots (771/3,719) (Chi-squared p=5.17×10^-182^) (**Fig 1D-E, S2B**). Within metastatic samples, 6.1% of PRAD-high spots co-localize with high stromal content (515/8,486) compared to only 0.1% of NEPC-high spots (2/2,334) (Fisher’s p=1.29×10^-52^) (**Fig S2C**). Recognizing that comparing primary mouse prostate tumors and metastatic human CRPC specimens has limitations, the relative depletion of stromal cells in association with NEPC in both settings raises the question of a potential functional link to lineage transition.

### ECM withdrawal initiates PRAD to NEPC lineage transition

Organoids are grown in Matrigel, an ECM preparation containing laminin (60%) and collagen IV (30%) that replicates the stroma-rich TME of PRAD. Because the NEPC transition is associated with depletion of stroma, we considered the possibility that Matrigel prevents the NEPC transition from occurring *in vitro*. To test this hypothesis, we removed Matrigel by growing TKOM organoids in suspension and monitored *Ascl1* expression as an early marker of NE lineage differentiation. *Ascl1* is not expressed in the luminal adenocarcinoma lineage but is essential for commitment to a neuroendocrine (NE) fate (*9*, *13*). Notably, *Ascl1* protein expression was detected by IHC in a subset of cells within individual organoids after 30 days of suspension culture (**Fig. 1F**). To explore the kinetics of *Ascl1* induction, we performed a time course experiment and documented induction of *Ascl1* mRNA by RT-PCR after 7 days in suspension, with increased levels after 14 days and reaching peak levels at 30 days (**Fig. S3A**). Of note, *Ascl1* scored as the top upregulated gene in suspension culture by bulk RNA sequencing (RNA-seq) among other NEPC-associated TFs, notably *Foxa2*, *Insm1*, and *Sox2* (**Fig. 1G, S3B**). Furthermore, gene signatures associated with NEPC were enriched in suspension culture while those associated with PRAD downregulated (**Fig. 1H-I**). Thus, removal of Matrigel from TKOM organoids is sufficient to activate a transcriptional program resembling NEPC.

To explore the dynamics of this phenotypic shift at a single cell level, we performed multiome (scRNA and scATAC) sequencing of TKOM organoids grown in Matrigel, or in suspension culture for 2 days or 20 days. *Ascl1* mRNA was detectable after 20 days but not in all cells (**Fig. S3C-D**), consistent with the IHC data, and closely matched with changes in chromatin accessibility at the Ascl1 locus (**Fig. S3E**). *Foxa2* and *Insm1* were also detectable at day 20 in a subfraction of the Ascl1+ cells (**Fig. S3F-G**).

### ECM-integrin signaling maintains PRAD lineage

While the suspension culture experiments implicate components of Matrigel as inhibitors of NE lineage transition in organoid culture, interpretation is confounded by the fact that prolonged growth of epithelial cells in suspension culture can enrich for other phenotypes such as acquisition of stem-like properties (*21*). The primary constituents of Matrigel, laminin and collagen IV, signal through distinct alpha integrin receptors but share a common beta receptor subunit, integrin beta-1 (ITGB1) (*22*). To ask if the induction of *Ascl1* seen in suspension culture is a consequence of impaired integrin signaling, we deleted *Itgb1* in TKOM organoids. Remarkably, *Itgb1* knockout replicated the mosaic pattern of ASCL1 protein expression observed in suspension culture but now in organoids grown in Matrigel (**Fig. 2A-B**). *Itgb1* loss also resulted in upregulation of *Foxa2* and *Insm1*, as well as the NE markers C*hga, Syp* and *Dll3* , together with downregulation of PRAD lineage markers such as *Hoxb13* and *Nkx3.1* (**Fig. 2C, S4A**). These findings establish that ECM-integrin signaling in prostate organoids is critical to maintain the PRAD lineage state and acts as a brake on the NEPC transition *in vitro*.

**Fig. 2.**
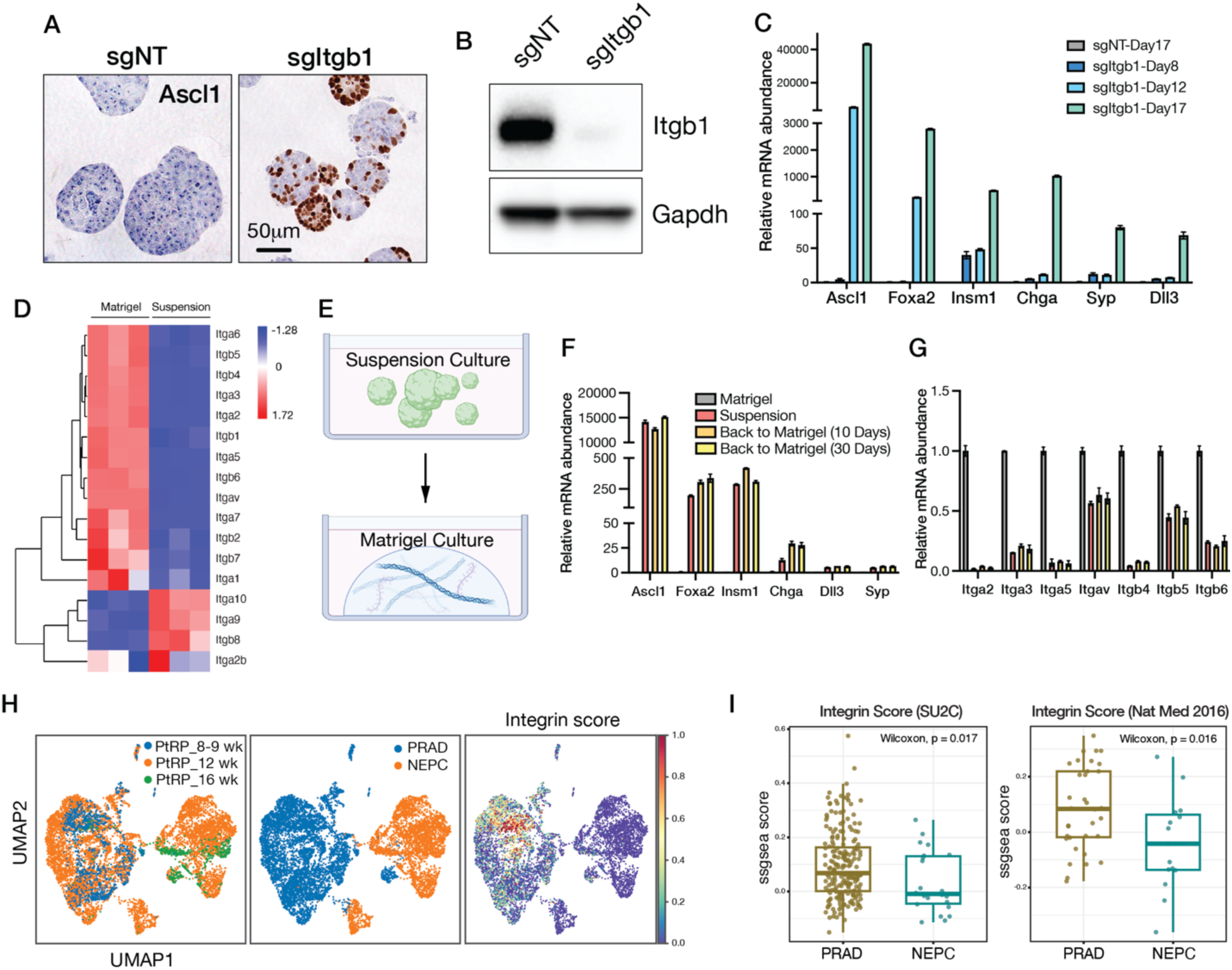
Loss of ECM-integrin signaling promotes PRAD to NEPC lineage transition. (A) ASCL1 IHC of TKOM organoids 17 days after electroporation with control sgRNA (sgNT) or sgRNA targeting Itgb1 (sgItgb1). Organoid fragmentation post Itgb1 knockout may result from weakened cell–cell adhesion and compromised junctional integrity. (B) Western blot confirming Itgb1 knockout in TKOM organoids. (C) Quantitative reverse-transcription PCR (qRT-PCR) analysis showing expression of NE markers at day 8, 12 and 17 following *Itgb1* deletion. Error bars represent ±SEM, n=3. (D) Heatmap showing differential expression of integrin genes in TKOM organoids cultured in Matrigel versus suspension. (E) Schematic diagram of the replating experiment. (F) qRT-PCR of NE markers in TKOM organoids cultured in Matrigel, suspension conditions and replating back to Matrigel for 10 or 30 days. Error bars represent ±SEM, n=3. (G) qRT-PCR showing the expression of multiple integrin genes in TKOM organoids cultured in Matrigel, suspension conditions and after replating back in Matrigel. Error bars represent ±SEM, n=3. (H) scRNA-seq analysis of PtRP (TKO) GEMMs showing reduced integrin gene expression score in NEPC compared to PRAD. (I) Box plot demonstrating reduced integrin gene expression score in NEPC (CRPC-NE) compared to PRAD (CRPC-adeno) in two published patient cohorts. The integrin gene expression score represents the mean expression of all integrin genes.

We next considered whether and how ECM-integrin signaling might be disrupted in the *in vivo* setting. To model disrupted ECM signaling, we examined gene expression changes seen after transfer of TKOM organoids to suspension culture and noticed that 7 alpha integrins (*Itga1,2,3,5,6,7,v*) and 5 beta integrins (*Itgb1,4,5,6,7*) were significantly downregulated (**Fig. 2D**). To determine if this level of downregulation has functional consequences, we replated suspension cells back into Matrigel to reactivate integrin signaling but found that NE marker expression remained stable at 10 and 30 days after replating (**Fig. 2E-F**). Thus, activation of the NE lineage program that follows loss of ECM-integrin signaling cannot be reversed by reengagement with ECM, likely due to sustained downregulation of integrin expression (**Fig. 2G**). Using an integrin score that quantifies expression of multiple integrin genes, we also observed significantly reduced expression in NEPC versus PRAD tumor cells in PtRP mice (**Fig. 2H**) and in human CRPC cohorts with PRAD and NEPC patients (**Fig. 2I**) (*12*, *23*).

To summarize, ECM-integrin signaling restricts the PRAD to NEPC lineage transition in organoid culture. Once this signal is disrupted, loss of integrin expression by tumor cells precludes reversion back to a PRAD lineage state, even if abundant ECM is present. These findings support a model where the abundant stromal population associated with the PRAD state *in vivo* restrains lineage plasticity. Consistent with this model, NEPC emerges in regions with a relative paucity of stromal cells.

### YAP1/TAZ/TEAD suppresses Ascl1 induction and NE lineage transition

To elucidate which pathways downstream of integrin engagement suppress the induction of Ascl1 expression, we first examined canonical kinase signaling events associated with integrin activity (*24*). pFAK, pSRC and pAKT were attenuated when cells were cultured in suspension or following *Itgb1* knockout, confirming that the circuitry of integrin signaling that is well defined in other cell types is conserved in prostate organoid culture (**Fig**. **3A-B**). We also examined the status of the YAP1/TAZ/TEAD (HIPPO) pathway (hereafter called YAP/TEAD), a target of integrin signaling linked to mechanical sensing (ECM stiffness) (*25*). Loss of integrin engagement resulted in reduced YAP/TEAD signaling, as measured by increased pYAP1 (a substrate for LATS1/2 kinase) and reduced expression of the YAP/TEAD target gene CYR61 (**Fig**. **3A-B**). To determine which, if any, of these signaling events are involved in *Ascl1* upregulation, we used a panel of small molecule inhibitors targeting FAK, SRC, PI3K, MEK or the YAP/TEAD complex to test if any might reproduce the *Ascl1* induction phenotype seen with *Itgb1* KO. Of the kinase inhibitors, only the SRC family inhibitor dasatinib gave a reproducible (∼30-fold) increase in *Ascl1* expression. More striking, however, was a ∼300-fold increase in *Ascl1* seen with IAG933, which selectively disrupts the YAP/TEAD protein-protein interaction (**Fig. 3C**) (*26*). IAG933 (hereafter called YAPi) also blocked expression of the YAP/TEAD target gene *Ctgf* and *Cyr61* (∼90% decrease), as did dasatinib (∼50% decrease), consistent with the known role of SRC kinases in activating YAP/TEAD through LATS1/2 inhibition (**Fig. 3D**) (*27*).

**Fig 3.**
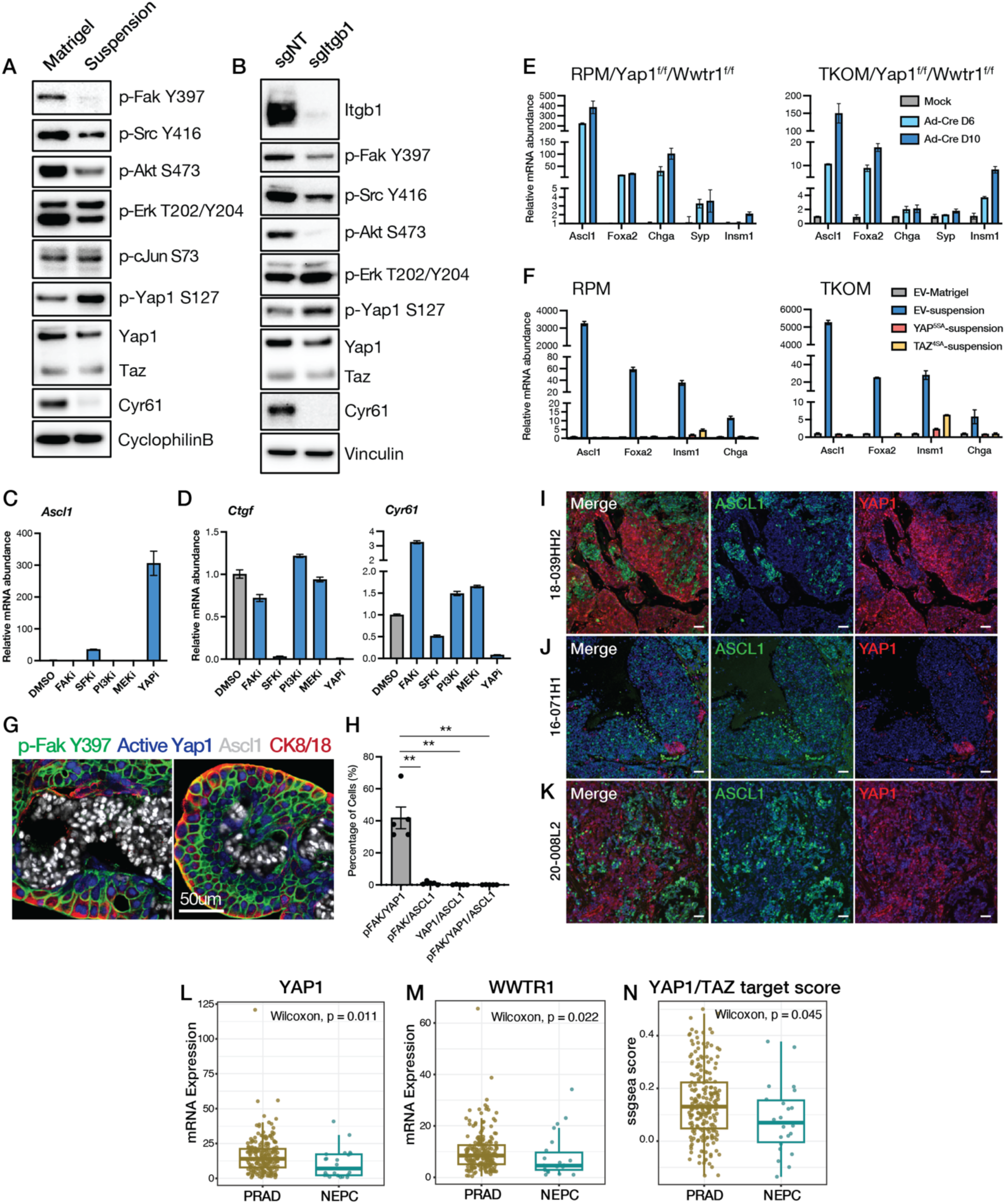
YAP/TEAD activation blocks Ascl1 induction. (A-B) Western blot analysis of representative kinases, substrates and target genes downstream of integrin downstream in TKOM organoids in Matrigel or after one day in suspension culture (A) or two days after Itgb1 knockout (B). (C) qRT-PCR for *Ascl1* in TKOM organoids after treatment with pharmacologic inhibitors of the indicated targets. FAKi: 5uM PF573228; SFKi: 5uM Dasatinib; PI3Ki: 2uM GDC0941; MEKi: 1uM AZD6244; YAPi: 1uM IAG933. Error bars, ±SEM, n=3. (D) qRT-PCR of the YAP1/TAZ target genes *Ctgf* and *Cyr61* in TKOM organoids after treating with the indicated inhibitors. Error bars, ±SEM, n=3. (E) qRT-PCR of NE genes in RPM/*Yap1^f/f^/Wwtr^f/f^* and TKOM/*Yap1^f/f^/Wwtr^f/f^*organoids 6 and 10 days following Adeno-Cre (Ad-Cre) infection. Error bars, ±SEM, n=3. (F) qRT-PCR of NE markers in RPM and TKOM organoids overexpressing EV, constitutive YAP5SA or TAZ4SA, in cultured in Matrigel or in suspension for 14 days. Error bars, ±SEM, n=3. (G-H) Multiplex IF of phospho-FAKY397, active-YAP1, ASCL1 and CK8/18 of tumor sections from 9-week-old TKO GEMM (G) with quantification (H). Error bars, ±SEM, n=3, **p<0.005. (I-K) Co-immunofluorescence imaging of three independent human CRPC clinical specimens containing admixed neuroendocrine prostate cancer (NEPC) and prostatic adenocarcinoma (PRAD) components demonstrates mutually exclusive expression of YAP1 (red) and ASCL1 (green). Nuclei were counterstained with DAPI (blue). Scale bars, 50 μm. 18-039HH2 is from vertebral bone metastasis. 16-071H1 is liver metastasis. 20-008L2 is from retroperitoneal lymph node metastasis. (L-N) Box plot demonstrating reduced expression of YAP1 (L), WWTR1 (M) and decreased YAP/TAZ target score (N) in NEPC compared to PRAD tumors from the SU2C cohort.

To build on the pharmacologic evidence implicating YAP/TEAD as a critical regulator of *Ascl1* induction, we turned to genetic experiments using gain and loss of function approaches. First, we generated RPM and TKOM mouse prostate organoids in a *Yap1^fl/fl^*/*Wwtr1^fl/fl^* background (the *Wwtr1* gene encodes TAZ) (*28*) to ask whether deletion of these TEAD co-activators would activate NE gene expression. We observed >100-fold induction of *Ascl1* and upregulation of other NE genes 6-10 days after co-deletion of *Yap1*/*Wwtr1* by adenoviral Cre infection, as well as the expected loss of YAP/TEAD target gene expression (*Cyr61*, *Ctgf*, *Axl, Ptpn14*) **(Fig. 3E, S4B-C)**. *Yap1*/*Wwtr1* co-deletion also resulted in downregulation of several alpha (*Itga2*, *Itga3, Itga5*, *Itga7, Itgav*) and beta integrin genes (*Itgb3, Itgb4, Itgb5, Itgb6, Itgb7, Itgb8*) (**Fig. S4D)**, mirroring the loss of integrin expression seen earlier when *Yap1*/*Wwtr1* intact organoids were placed in suspension culture (**Fig. 2D**). Thus, YAP/TEAD is a key regulator of integrin expression in this context. Conversely, expression of a constitutively active mutant of either YAP (YAP^5SA^) or TAZ (TAZ^4SA^), both of which are resistant to inactivation by LATS1/2 phosphorylation, blocked the induction of *Ascl1* mRNA and other NE markers (*Foxa2*, *Insm1*, *Chga*) when RPM and TKOM organoids were placed in suspension culture, while maintaining YAP/TAZ target gene expression (*Cyr61*, *Ctgf*, *Axl, Ptpn14*) and integrin expression (**Fig. 3F, S4E-F**).

Collectively, the organoid experiments provide evidence that ECM-integrin engagement reinforces PRAD lineage identity through YAP/TEAD activation. To determine whether ECM-integrin engagement is also linked to YAP/TEAD activity *in vivo*, we performed multiplex immunofluorescence (IF) on prostate tissue sections from 9-week-old PtRP mice, when NEPC first starts to emerge from PRAD. Using phosphoFAK (pFAK) as a readout for ECM-integrin engagement, we found that loss of pFAK in tumor cells was highly correlated with reduced nuclear YAP1 staining and gain of ASCL1 expression as measured by multiplex immunofluorescence (*p*<0.005) (**Fig. 3G-H**). To determine whether the reciprocal association of YAP1 activity and ASCL1 expression is also present in human NEPC, we performed a similar analysis of metastatic samples from three CRPC patients with mixed lineage PRAD/NEPC histology. As in mouse, nuclear YAP1 activity and ASCL1 expression were mutually exclusive (**Fig 3I-K**). Also, YAP/TEAD activity in human CRPC cohorts (via RNA signatures) was significantly reduced in NEPC compared to PRAD (**Fig. 3L-N**), providing further evidence for this reciprocal association and consistent with scRNA-seq data from a smaller CRPC cohort (*29*). In sum, data from organoids, GEMMs and human CRPC supports a model whereby ECM-integrin engagement maintains the PRAD lineage through sustained YAP/TEAD signaling.

### LATS inhibition impairs acquisition and maintenance of neuroendocrine state

Earlier we showed that the PRAD to NEPC lineage transition following loss of ECM-integrin engagement cannot be reversed by re-exposure to ECM - likely due to loss of integrin expression (**Fig. 2E-G**). However, the fact that YAP1/TEAD activity is regulated by LATS1/2 kinase activity (YAP1 and TAZ phosphorylation by LATS1/2 results in cytoplasmic retention and inability to bind TEAD (*30*)) led us to ask whether the reduced YAP/TEAD activity caused by loss of ECM-integrin engagement could be restored through LATS kinase inhibition (**Fig. 4A**). Indeed, treatment of TKOM organoids with the LATS1/2 kinase inhibitor TRULI (*31*) (hereafter called LATSi) blocked YAP1 phosphorylation induced by suspension culture and restored expression of canonical YAP/TEAD target genes such as CYR61 (**Fig. 4B**). Continued treatment for 14 days prevented induction of *Ascl1* and other NE markers (*Foxa2*, *Insm1*, *Chga*) (**Fig. 4C, S5A**). Furthermore, induction of ASCL1 was reversed when LATSi was added after prolonged growth (30 days) in suspension culture (**Fig. 4D**).

**Fig. 4.**
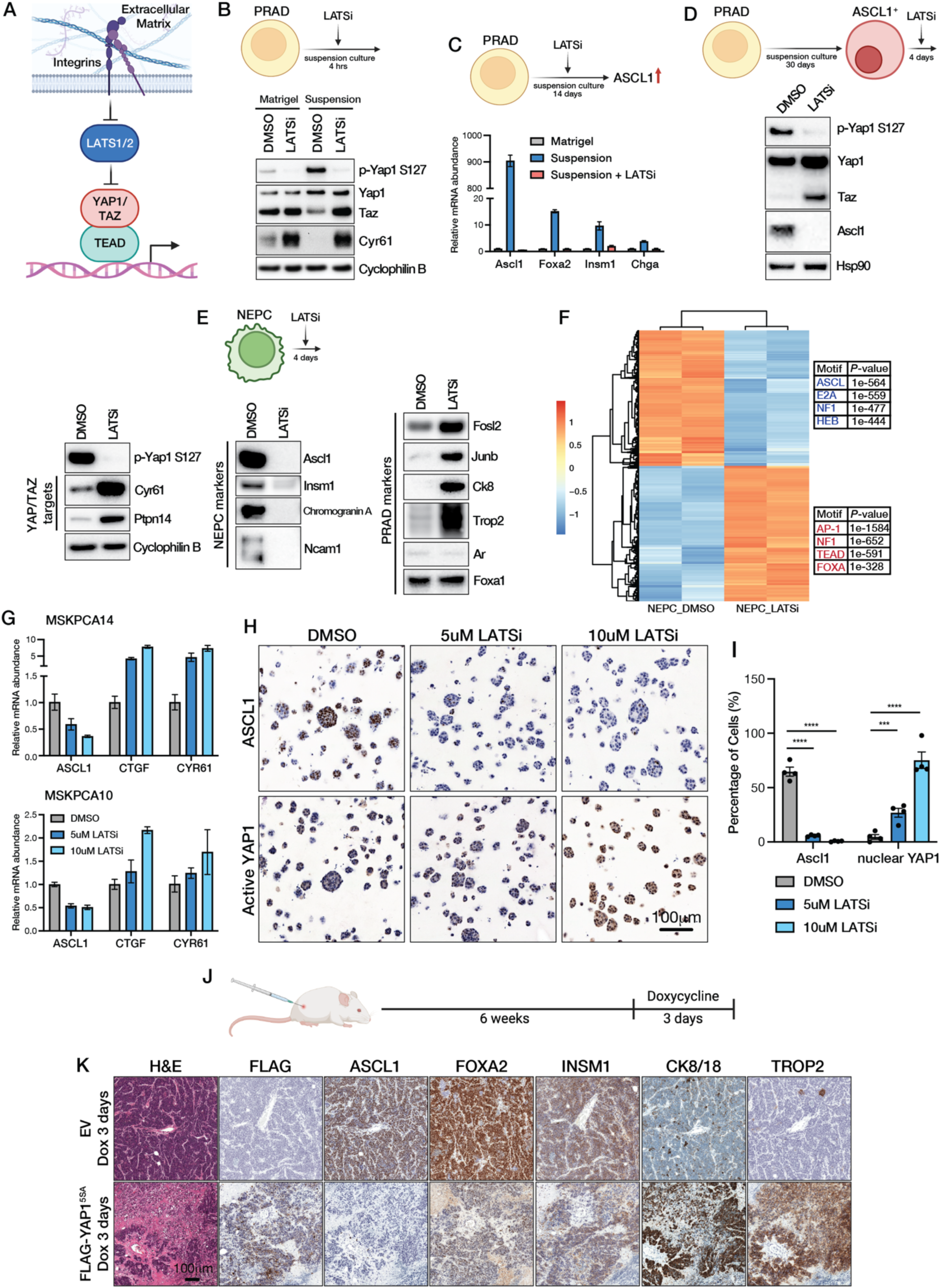
LATS inhibition restores YAP/TEAD activity and reverses NE lineage transition. (A) Cartoon showing ECM-Integrin inhibits LATS1/2 to activate YAP1/TAZ/TEAD. (B) Upper: Schematic of the experimental design. Lower: Western blot analysis of indicated proteins in TKOM organoids cultured in Matrigel or suspension, treated with LATSi (5uM TRULI) for 4 hours. (C) Upper: Schematic of the experimental design. Lower: qRT-PCR of NE genes (*Ascl1*, *Foxa2*, *Insm1*, and *Chga*) in TKOM organoids cultured in Matrigel, suspension, or suspension plus LATSi (5uM TRULI) for 14 days. Error bars represent ±SEM, n=3. (D) Upper: Schematic of the experimental design. Lower: Western blot analysis of indicated targets in RPM PRAD organoids cultured in suspension for 30 days followed by treatment with LATSi (5uM TRULI) for 4 days. (E) Upper: Schematic of the experimental design. Lower: Western blot analysis of indicated targets in RPM NE organoids following treatment with LATSi (5uM TRULI) for 4 days. (F) Unsupervised clustering of ATAC-seq peaks in RPM-NEPC organoids treated with DMSO or LATSi (5uM TRULI) for 8 days (left). Top enriched transcription factor motifs and associated *P* values for each cluster (right). (G) qRT-PCR analysis of ASCL1 and YAP1/TAZ targets (CTGF and CYR61) in MSKPCA14 and MSKPCA10 patient-derived organoids treated with DMSO, 5uM LATSi (TRULI), 10uM LATSi (TRULI) for 10 days. Error bars represent ±SEM, n=3. (H) ASCL1 and YAP1 IHC in MSKPCA14 patient-derived organoids treated with DMSO, 5uM LATSi (TRULI), or 10uM LATSi (TRULI) for 10 days. (I) Quantification of the percentage of ASCL1 positive cells and nuclear YAP1 positive cells. Error bars represent ±SEM, n=4, ***p<0.0005, ****p<0.0001. (J) Schematic showing the timeline of the subcutaneous transplantation experiment using RPM organoids. (K) Representative H&E images and IHC images of the indicated markers in RPM subcutaneous tumors overexpressing empty vector (EV) or YAP^5SA^, n=8 mice per group.

The above experiments establish that induction of ASCL1 upon transfer of PRAD organoids to suspension culture is reversible. To ask whether LATSi can reverse the fully established NEPC tumor phenotype that develops *in vivo*, we established tumoroids from ASCL1+ NEPC tumors that developed following orthotopic transplantation of RPM and TKO organoids (see methods for details) (**Fig S5B-E**). Again, LATSi blocked YAP1 phosphorylation and restored YAP/TEAD activation, as shown by induction of target genes such as CYR61 and PTPN14. Remarkably, after 4 days of treatment, these NEPC tumoroids had near complete loss of ASCL1, INSM1 and CHGA expression and gained expression of PRAD lineage markers such as CK8, TROP2 and the AP1 complex proteins FOSL2 and JUNB (**Fig. 4E**). This switch in lineage marker expression was accompanied by a change in chromatin landscape, with closure of peaks enriched for ASCL1 motifs and opening of peaks enriched for TEAD motifs (**Fig 4F, table S3)**. Globally, the newly open chromatin peaks map to canonical PRAD lineage genes whereas the closed chromatin peaks map to canonical NEPC lineage genes, consistent with enhancer reprogramming (**S5F, table S2**). Collectively, these data show that an established NEPC transition can be reverted back to a PRAD-like state by pharmacological restoration of YAP/TEAD activity.

Having shown that the PRAD to NEPC lineage transition can be blocked or reversed by LATSi in mouse tumoroid models, we asked whether this reversion extends to human NEPC. Using patient-derived organoids (PDOs) established from CRPC patients with NEPC (*32*, *33*), we found that LATSi activated YAP/TEAD target gene expression (*CYR61*, *CTGF*) in a concentration-dependent manner, confirming the same pathway is intact in human CRPC models (**Fig. 4G**). Treatment with LATSi robustly restored YAP1 nuclear protein expression, coupled with a reduction in the percentage of ASCL1 expressing cells from >60% to <5% (**Fig. 4H-I**), further credentialling LATS1/2 kinase as a potential therapeutic target to prevent or reverse PRAD to NEPC lineage plasticity.

Unfortunately, we were unable to test this hypothesis in mice using TRULI or the chemically related LATSi TDI-011536 (*34*) due to rapid *in vivo* clearance and full recovery of pYAP in prostate tissue 4 hours after drug administration (**Fig S5G**). We therefore turned to a genetic approach, using the previously described constitutively active YAP1^5SA^ allele but now in a doxycycline (Dox) inducible format such that we could enforce YAP activation in RPM organoids at various times during the PRAD to NEPC transition *in vivo*. As in organoid culture (**Fig 3F**), induction of YAP1^5SA^ expression 2 weeks after transplantation blocked the appearance of detectable ASCL1+ cells in the PRAD tumors that emerged 21 days later, whereas control tumors transduced with the Dox empty vector control (EV) had abundant regions of ASCL1+ NEPC (**Fig S5H-J**). We conclude that sustained YAP/TEAD activation can prevent PRAD to NEPC lineage plasticity *in vivo*.

To determine if YAP activation can restore PRAD lineage marker expression in tumors that have already undergone the NEPC transition, we delayed YAP1^5SA^ induction until 6 weeks after transplantation, when large sections of harvested tumors have fully transitioned to ASCL1^+^, FOXA2^+^, INSM1^+^ NEPC (**Fig 4J-K**, EV_Dox). Remarkably, ASCL1, FOXA2 and INSM1 expression was reversed after 3 days of YAP1^5SA^ induction and expression of PRAD lineage marker (TROP2, CK8/18) was restored (**Fig 4K**, FLAG-YAP1^5SA^_Dox). Taken together, the organoid and *in vivo* data provide proof of concept that PRAD to NEPC lineage plasticity can be reversed, at least during the early stages of the transition.

### Cooperativity of YAP1/TEAD, NOTCH and AR signaling in PRAD lineage maintenance

The above data provide evidence that disruption of YAP/TEAD activity (through suspension culture, *Itgb1* deletion or pharmacologic blockade of YAP/TEAD binding) promotes NE lineage transition. However, *in situ* analysis of ASCL1 protein expression reveals a mosaic pattern with all three methods of YAP/TEAD perturbation (**Figs. 1F**, **2A**, **5C**) suggesting that other regulatory pathways may play a role. In addition, the mRNA levels of *Ascl1* and other NE markers (*Foxa2*, *Insm1*, *Ncam*) achieved in organoids following YAP/TEAD perturbation do not reach the level seen in NEPC tumors that develop in organoids transplanted *in vivo* (**Fig. S6A**).

**Fig. 5.**
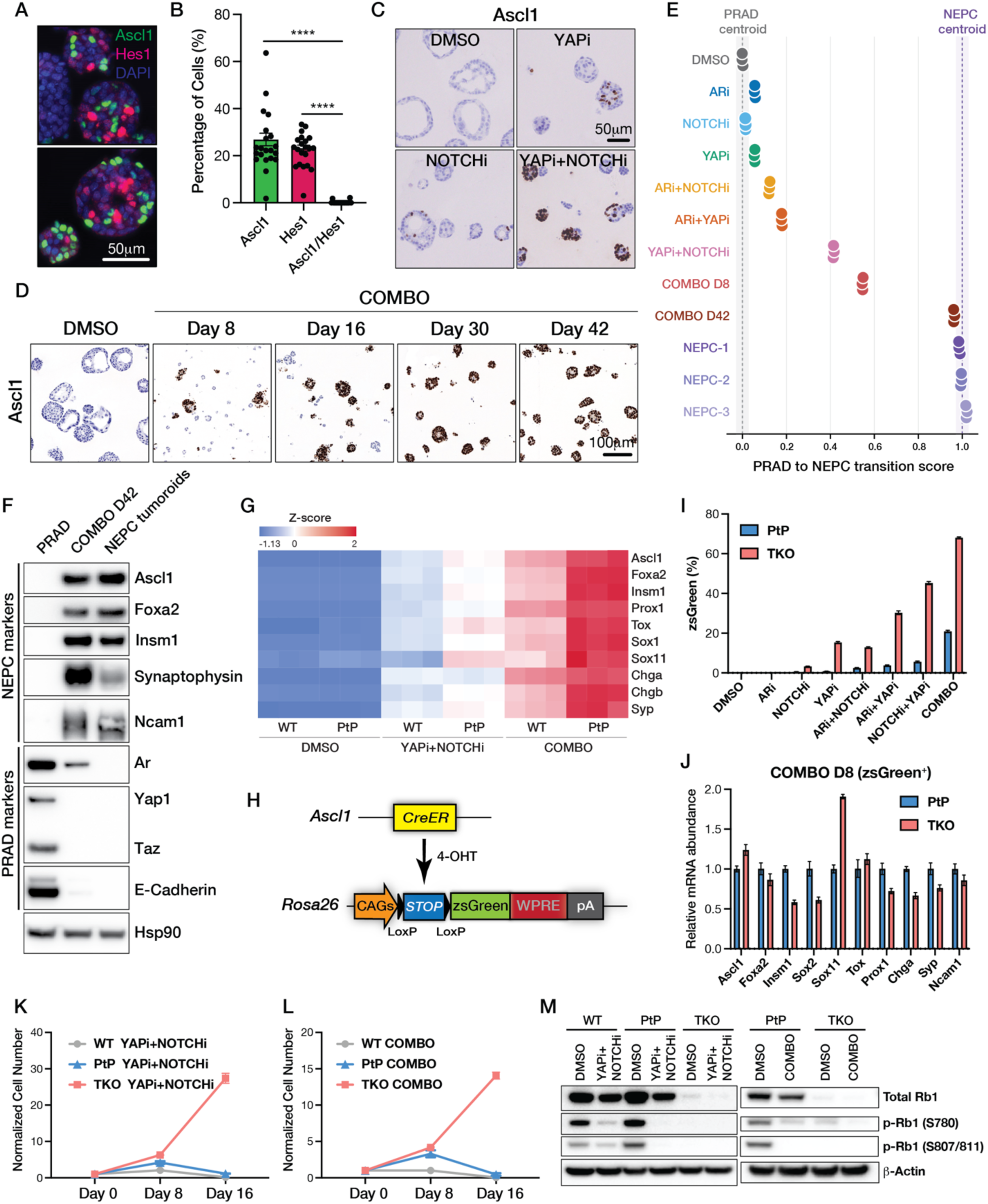
Reconstitution of PRAD to NEPC lineage transition *in vitro*. (A and B) Multiplex IF staining for ASCL1, HES1 and DAPI in TKOM organoids cultured in suspension (A) and quantified (B). Error bars, ±SEM, n=20, ****p<0.0001. (C) ASCL1 IHC in TKO PRAD organoids with DMSO, YAPi (1uM IAG933), NOTCHi (10uM DAPI) or a combination of YAPi and NOTCHi for eight days. The scale bar represents 50um. (D) ASCL1 IHC in TKO PRAD organoids treated with COMBO treatment over the indicated time course. The scale bar represents 100um. (E) PRAD-to-NEPC transition scores derived from bulk RNA-seq profiles of PRAD organoids treated under the indicated conditions, together with three established NEPC models. The PRAD and NEPC reference states were defined by the mean expression profiles of the three DMSO samples and all nine NEPC samples, respectively. Each sample was projected onto the transcriptional axis connecting the PRAD and NEPC reference centroids, with scores scaled such that the PRAD centroid equals 0 and the NEPC centroid equals 1. Each circle represents an independent biological replicate. Dashed lines indicate the PRAD and NEPC reference centroids. (F) Western blot of NEPC and PRAD lineage markers in TKO PRAD organoids, TKO PRAD organoids treated with COMBO for 42 days (COMBO D42), and NEPC tumoroids. (G) Heatmap showing bulk RNA-seq expression of NE lineage-associated genes in WT and PtP (*Pten^-/-^*, *Trp53*^-/-^) organoids treated under the indicated conditions. Expression levels are shown as Z-score. (H) Schematic of the Ascl1 reporter system. (I) Flow cytometry quantification of the percentage of zsGreen-positive cells in PtP and TKO (*Pten^-/-^*, *Trp53*^-/-^, *Rb1^-/-^*) organoids treated under the indicated conditions for 8 days. Error bars, ±SEM, n=3. (J) qRT-PCR analysis of NE marker gene expression in zsGreen-positive cells sorted from PtP and TKO organoids following COMBO treatment for 8 days. Error bars, ±SEM, n=3. (K-L) Growth curves of WT, PTP, and TKO organoids under YAPi and NOTCHi (K) or COMBO (L) treatment. Cell numbers were normalized to Day 0 and measured at the indicated time points. Data are shown as mean ± SEM, n=3. (M) Western blot analysis of total RB1 and phosphorylated RB1 (p-RB1 S780 and S807/811) in WT, PTP, and TKO organoids treated with DMSO, DOUBLE, or COMBO. β-Actin serves as a loading control. The duration of the treatment is 24h.

The mosaic pattern of ASCL1 expression is reminiscent of segmental patterns of protein expression seen in normal tissue development, often a consequence of cell-cell communication through signaling pathways such as NOTCH. Furthermore, NOTCH is known to suppress NE differentiation across various tissues, including in models of NEPC and ASCL1+ small cell lung cancer (*35–37*). To determine if NOTCH plays a similar role in prostate organoids, we first performed dual IF staining of ASCL1 and the NOTCH target HES1 in TKOM organoids grown in suspension culture. ASCL1 and HES1 were robustly expressed in a mutually exclusive pattern in all organoids examined (**Fig. 5A-B**). To test the hypothesis that NOTCH activation prevents *Ascl1* induction in the TKO model, we treated organoids with the gamma secretase inhibitor DAPT (hereafter called NOTCHi) (*38*) alone or in combination with YAPi for 8 days. NOTCHi alone resulted in a mosaic pattern of ASCL1 expression, reminiscent of that seen with YAPi albeit less pronounced. However, combination treatment with NOTCHi and YAPi resulted in intense ASCL1 expression in nearly all the cells within each organoid (**Fig. 5C**), indicative of cooperativity between NOTCH and YAP/TEAD activation in suppressing ASCL1. Since ARPI therapy typically precedes the emergence of NEPC, we also measured the consequences of AR inhibition. Addition of enzalutamide plus DHT withdrawal (hereafter called ARi) to NOTCHi and YAPi treatment (hereafter called COMBO) further enhanced *Ascl1* induction (**Fig. S6B-D**) coupled with a progressive increase in the percentage of organoids that now uniformly express ASCL1 over 30-42 days **(Fig. 5D)**.

### Complete reconstitution of PRAD to NEPC lineage transition *in vitro*

The robust and homogeneous pattern of ASCL1 expression seen after 6-7 weeks of COMBO treatment suggests PRAD organoids may undergo progressive reprogramming to a complete NEPC lineage state. To address this possibility, we compared the transcriptomes of PRAD organoids treated with vehicle control (DMSO), single inhibitors (YAPi, NOTCHi or ARi), doublet combinations (YAPi /NOTCHi, YAPi/ARi, NOTCHi/ARi) and COMBO in a time course experiment. Using the time course of ASCL1 expression as a guide (**Fig. 5D**), we harvested organoids at 8 days to capture an intermediate state of lineage reprogramming and at 42 days to represent the final NE lineage state achieved *in vitro*. As an additional control, we included three independently derived NEPC tumoroids (NEPC1-3) to compare the NE state generated by *in vitro* reprogramming with the fully reprogramed NEPC state that develops *in vivo*.

Projecting the samples along a transition axis (0.0-1.0) defined as the difference between reference profiles for PRAD (defined by DMSO) and NEPC (defined from NE tumoroids) (**table S4**), we found that each single treatment and both ARi doublet treatments had transition scores modestly increases relative to the DMSO control (**Fig. 5E**). In stark contrast, organoids treated for 42 days with COMBO had transition scores approaching those of the three NE tumoroids, suggesting that sustained COMBO treatment of PRAD organoids *in vitro* replicates the full NEPC transition seen *in vivo*. The 8-day timepoint for the YAPi/NOTCHi doublet and COMBO treatments were similarly revealing because both have intermediate transition scores (0.4-0.6), suggestive of a transition state between PRAD and NEPC. To verify that the extremes of the transition scores do indeed reflect the PRAD and NEPC states respectively, we generated heatmaps using previously reported PRAD and NEPC mRNA signatures (from the PtRP GEMM) (*17*) across all treatment conditions and timepoints. Indeed, the NE signature is high in NE tumoroids and COMBO but absent in DMSO and all single inhibitor (and ARi combo) treatments. Conversely, the PRAD signature is high in DMSO and all single inhibitor (and ARi combo) treatments but low in NE tumoroids and COMBO (**Fig. S6F-G**). As further confirmation, expression of NE markers at the protein level (ASCL1, FOXA2, INSM1) after 42 days of COMBO are comparable to those seen in NE tumoroids, as well as loss of PRAD markers such as E-cadherin (**Fig. 5F**). In sum, these data establish an organoid platform for complete reprogramming from PRAD to NEPC through combined and sustained inhibition of YAP, NOTCH and AR.

Because *RB1* loss is a critical molecular determinant of the PRAD to NEPC transition *in vivo* (*9*), we presumed that *RB1* loss might also be required for induction of the NE lineage program in organoids treated with YAP, NOTCH and AR inhibitors. However, to our surprise, we observed induction of canonical NE lineage genes (*Ascl1*, *Foxa2*, *Insm1*, etc.) in *Rb^+/+^*, *Pten^-/-^*, *Tp53^-/-^* (PtP) as well as wildtype (WT) prostate organoids after 8 days of COMBO treatment, albeit not to the same magnitude as in *Rb^-/-^*, *Pten^-/-^*, *Tp53^-/-^* (TKO) organoids (**Fig 5G, S7A-B**). Because these measurements were derived from bulk RNA-seq data, the genotype-specific differences in induction of NE lineage genes might be due to differences in efficiency rather than magnitude of induction. To distinguish between these two possibilities, we derived organoids carrying an *Ascl1* reporter (*39*, *40*) to track induction on a cell-by-cell basis using zsGreen fluorescence as a readout (**Fig 5H, see methods**). This analysis revealed that *Ascl1* was induced in a much larger fraction of TKO cells (∼70%) compared to PtP cells (∼20%) but the magnitude of *Ascl1* (and other NE lineage transcripts) across RB^+/+^ and Rb^-/-^ genotypes was comparable (**Fig 5I-J**). Induction of the NE program in RB wildtype cells was also seen in human prostate cancer cells. Modest *ASCL1* induction was also observed following COMBO treatment in LNCaP cells (RB wildtype) but this induction was more potent in *RB-* and *TP53-*deleted LNCaP cells (**Fig S7C**). To summarize, the NE lineage program can be induced in all prostate cells by COMBO treatment regardless of genotype, but the efficiency of that induction is greatly enhanced by RB loss.

To investigate why RB loss is required for cells to transition fully to NEPC but not for induction of the NE transcriptional program, we tracked the growth of WT, PtP and TKO organoids treated with YAPi + NOTCHi or COMBO for 16 days. Interestingly, WT and PtP organoid growth slowed dramatically, with absence of Ki67 expression in *Ascl1*^+^ cells, where TKO cells proliferated throughout the treatment period (**Fig 5K-L, S7D-E**). Of note, growth arrest coincided with absence of RB phosphorylation within hours of treatment, evidence that COMBO induces a G1/S cell cycle checkpoint (**Fig 5M**). We conclude that YAPi + NOTCHi + ARi can induce a NE lineage specification program in all prostate cells, regardless of RB status, but only those with RB1 loss can progress to NEPC due to induction of a growth arrest program in cells with intact RB1.

### FOXA1 is essential for the PRAD to NEPC lineage transition

To explore the specific molecular events responsible for the PRAD to NEPC lineage transition in the context of *Rb1* loss (TKO organoids), we examined the YAP/TEAD axis, using CUT&RUN to map the chromatin binding sites of YAP1 and TEAD1 across the PRAD and NEPC lineages. Focusing initially on TEAD1 peaks, we observed a remarkable re-localization of binding sites when comparing the PRAD state to the NEPC state (**Fig. 6A, table S3**). In PRAD, genes and pathways enriched at the sites of TEAD peaks include actin filament organization, epithelial morphogenesis, and cell substrate adhesion, all consistent with an adenocarcinoma lineage (exemplified by *Tacstd2*). Conversely, genes and pathways enriched at the sites of the redistributed TEAD peaks in the NEPC state include regulation of neurogenesis, forebrain development and synapse pathways (exemplified by *Ascl1*) (**Fig. S8A-D**).

**Fig. 6.**
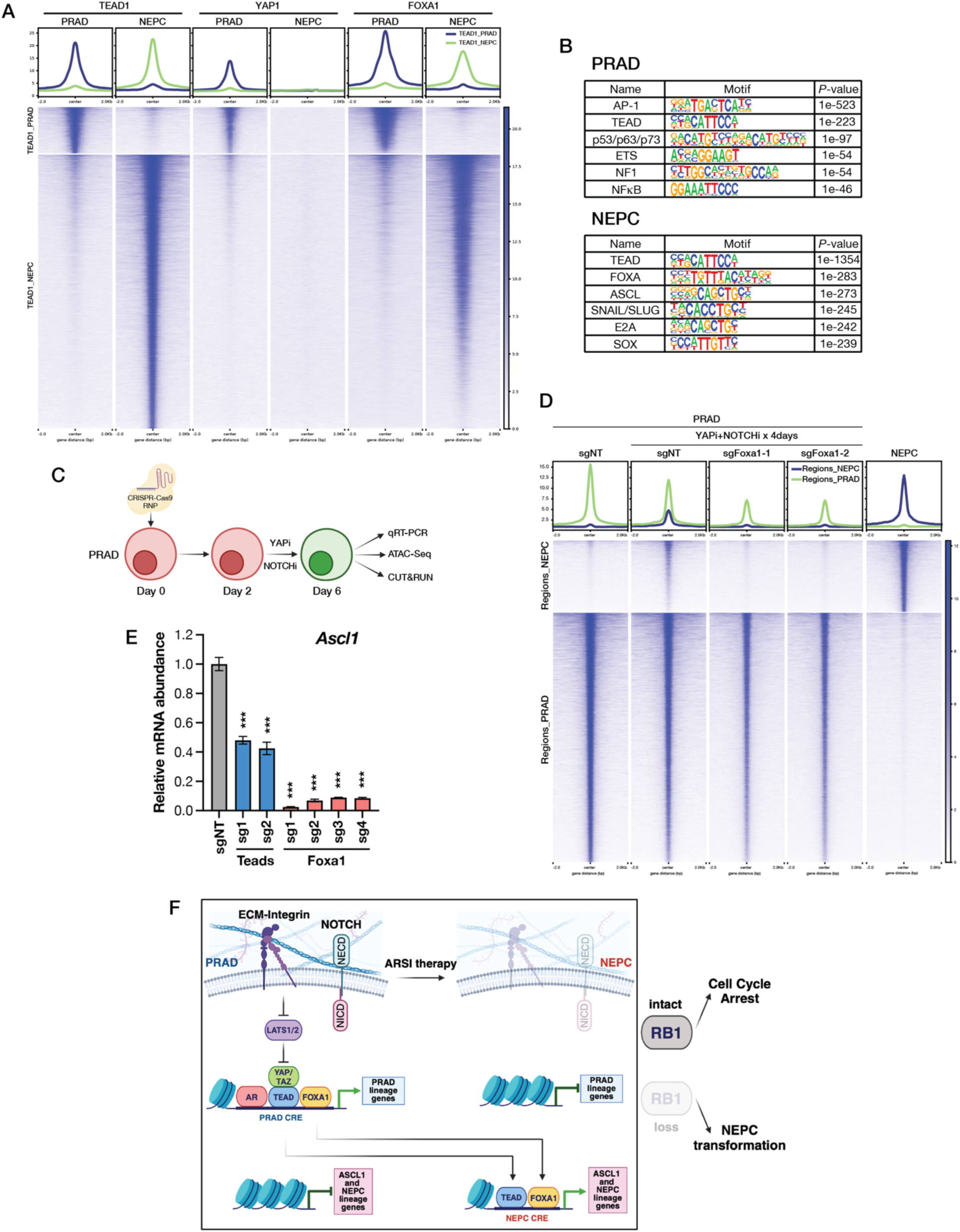
FOXA1 is essential for the PRAD to NEPC lineage transition. (A) CUT&RUN profiles of TEAD1, YAP1 and FOXA1 in TKO PRAD and NEPC organoids. Clusters were defined based on differential TEAD1 binding between PRAD and NEPC. (B) Top enriched transcription factor binding motifs identified by HOMER known-motif analysis of TEAD1 CUT&RUN peaks in TKO PRAD and NEPC organoids. (C) Schematic illustrating the experimental design used to assess the role of Teads and Foxa1 in driving the PRAD-to-NEPC transition. (D) ATAC-seq peak heatmaps showing chromatin accessibility across PRAD and NEPC-specific regions across the indicated organoid lines and conditions. Differential peaks were identified following depth normalization of ATAC-seq signal. (E) qRT-PCR for *Ascl1* in TKO PRAD organoids treated with YAPi (1uM IAG933) and NOTCHi (10uM DAPT) following knockout of the indicated TFs. Error bars, ±SEM, n=3, ***p<0.001. (F) Model illustrating how YAP/TEAD, NOTCH, AR, and RB1 regulate the PRAD to NEPC lineage transition. ECM–integrin signaling activates YAP/TEAD through inhibition of LATS1/2 to maintain PRAD lineage identity, in cooperation with NOTCH and AR signaling. Combined inhibition of YAP/TEAD, NOTCH, and AR reprograms PRAD cells into the NEPC lineage. Early during this transition, FOXA1 and TEAD redistribute from PRAD cis-regulatory elements (CREs) to NEPC CREs, which leads to opening of new chromatin, activation of *ASCL1* expression, and commitment to the NE lineage. In presence of functional RB1, induction of the NE lineage program is accompanied by cell cycle arrest, whereas RB1 loss permits continued proliferation that results in full NEPC transformation.

YAP1 peaks in the PRAD state overlapped precisely with TEAD1 peaks, as expected, since these two proteins form a complex but were absent in the NEPC state because LATS kinase activation disrupts the YAP/TEAD complex (**Fig. 6A**). The fact that robust TEAD peaks are now detected at the enhancers of NEPC-specific genes in the absence of YAP raises the question whether TEAD binding plays a role in their expression. To address this, we performed CRISPR experiments targeting TEAD1, the most abundantly expressed of the four TEAD family members in the NEPC state and achieved robust knockdown of TEAD1 expression, albeit with some residual TEAD expression detected using a pan TEAD antibody, presumably from other TEAD family members (**Fig. S8E**). Despite this caveat, *Ascl1*, *Chga* and *Ncam1* mRNA and protein were reduced by ∼50% following TEAD1 knockdown (**Fig S8F**), evidence that the re-localized TEAD peaks play a role in maintenance of NE-specific gene expression despite the absence of YAP and TAZ (**Fig 5F**). Because TEADs are DNA binding proteins without intrinsic coactivation function, it will be of interest to determine whether another (perhaps NE-specific) coactivator is recruited to perform this function.

To further investigate the biology underlying the NEPC-specific TEAD1 peaks, we performed motif analysis. TEAD binding motifs were enriched in both PRAD and NEPC lineages, but there were clear lineage-specific differences in co-enriched motifs, such as FOXA and ASCL1 binding sites in NEPC (**Fig. 6B**). The fact that FOXA motifs are co-enriched at the NEPC-specific TEAD binding sites raises the possibility that FOXA1, which is abundantly expressed in both PRAD and NEPC states, may play a role in the lineage transition. To explore this, we first performed CUT&RUN and confirmed that FOXA1 peaks are indeed present at the NEPC-specific TEAD binding sites and, like TEAD, re-localize from PRAD to NEPC specific enhancers (**Fig. 6A**). Integrative analysis of ATAC, RNA-seq and CUT&RUN data confirmed concordant binding of FOXA1 and TEAD during the PRAD to NEPC transition (**Fig S8G-I)**.

Because FOXA1 is a pioneer factor, we postulated that FOXA1 mobilization might be an early step in the NEPC transition, catalyzing the opening of closed chromatin at NEPC specific enhancer regions and thereby enabling binding of NEPC lineage TFs. To test this hypothesis, we treated the PRAD cells with DMSO versus YAPi + NOTCHi (+/- CRISPR deletion of *Foxa1*) to assess changes in open chromatin and to determine whether FOXA1 is required for the NE transition (**Fig. 6C**). ATAC-seq experiments revealed a global increase in open chromatin at NE-specific loci after 4 days that was diminished in FOXA1-deleted cells (**Fig. 6D, table S3**), consistent with the known role of FOXA1 as a pioneer factor. *Foxa1* deletion also abrogated *Ascl1* induction, consistent with the absence of FOXA1 peaks at the *Ascl1* enhancer following knockdown (**Fig. 6E, S9A**). *Foxa1* deletion also blocked the appearance of new TEAD peaks (**Fig S9B-C**), indicating that FOXA1 mobilization likely precedes TEAD mobilization during lineage transition. Like FOXA1, TEAD also plays a role in Ascl1 induction during the early stages of the NE transition as revealed by a ∼50% decrease following pan-*Tead* knockdown (**Fig. 6E**). In contrast, deletion of either *Foxa2* or *Insm1* had no effect (**Fig S9D-G)**, consistent with the fact that both genes are expressed later during the transition. Interestingly, *Insm1* is required for induction of *Chga* and *Syp*, suggestive of a distinct role in NE TF circuitry, potentially downstream from ASCL1. Together, these findings establish that FOXA1, likely through its role as a pioneer factor, is a key upstream regulator of *Ascl1* induction during the PRAD to NEPC transition.

## Discussion

Next generation ARPIs extend prostate cancer survival through increased selective pressure on AR. Despite this clinical success, tumor cells can escape AR dependence by transitioning from AR-dependent luminal cells to AR-independent NEPC cells (lineage plasticity). Here we show that the YAP/TEAD pathway plays a key role in maintenance of the PRAD lineage state, preventing tumor cells from undergoing the PRAD to NEPC lineage switch. Prior work on HIPPO pathway signaling, initially in flies and subsequently in mammalian cells, has established a critical role of YAP/TEAD in growth regulation (*30*, *41–43*). In the context of tumor cells, the role of YAP/TEAD in growth regulation is more nuanced. Studies with pharmacologic inhibitors reveal YAP/TEAD dependency in cancers with HIPPO pathway mutations (such as mesotheliomas with NF2 loss) but not more broadly, even in tumors with robust YAP/TEAD activation (*26*, *44*). Here we provide evidence here that YAP/TEAD functions as a gatekeeper, preventing prostate epithelial tumor cells at risk for lineage plasticity from undergoing a lineage transition.

Using PRAD as a model, we find ECM engagement is the primary mechanism of YAP/TEAD activation, consistent with prior evidence linking YAP/TEAD to integrins and mechanical stress (*25*). Once PRAD tumor cells lose contact with ECM (by experimental withdrawal in organoid culture or stochastically during *in vivo* tumor expansion), YAP/TEAD activity is acutely switched off, followed by induction of ASCL1 and an eventual transition to NEPC. Importantly, this switch cannot be easily reversed by reengagement with ECM because integrin expression (which itself is YAP/TEAD dependent) is lost. Thus, tumor cells poised for lineage transition (i.e., those with RB loss) enter a point of no return once the YAP/TEAD signal is switched off. Of note, YAP/TEAD has also been implicated in other lineage transitions such as reversion to a fetal-like state in a colorectal cancer model (*45*) and an epithelial mesenchymal-like state in a lung cancer model (*46*). Importantly, in these examples YAP/TEAD activity is oncogene-driven and promotes plasticity. In contrast, our work shows the ECM/integrin-driven YAP/TEAD activation restrains plasticity in PRAD cells primed for lineage transition due to *RB1* and *TP53* loss.

Interestingly, NE lineage transition following loss of ECM-integrin engagement occurs in a mosaic pattern, suggesting cell-cell interactions provide an additional brake on the PRAD to NEPC lineage independent of YAP/TEAD. Recognizing the critical role of NOTCH in cell-cell interactions, we explored NOTCH signaling in our organoid model through gamma secretase inhibition and found remarkable cooperativity of NOTCH and YAP/TEAD in maintenance of the PRAD state, consistent with evidence that NOTCH can suppress NE lineage transitions in prostate and lung cancer models (*35*, *36*).

We also find that ARPI therapy, the therapeutic intervention that initiates the NEPC transition in CRPC patients, contributes to the PRAD to NEPC transition, particularly when YAP/TEAD or NOTCH signaling (or both) have already been impaired. The mechanism is cell autonomous because the NE lineage transition can be fully elicited in organoid cultures that only contain tumor cells. A plausible explanation is that AR plays a role in maintenance of luminal lineage identity, even in CRPC. That said, non-cell autonomous mechanisms may also be at play. Intriguingly, ECM production by stromal cells (e.g., collagen) in the normal mouse prostate is AR-regulated (*47*). One consequence of systemic ARPI therapy could be reduced YAP/TEAD activation in tumor cells due to reduced ECM production by AR-positive stromal cells, thereby lowering the threshold for NEPC transition to occur.

Our ability to fully recapitulate the PRAD to NEPC transition *in vitro* provided an opportunity to investigate the molecular details of this dynamic process at a level of resolution and efficiency not easily achievable *in vivo*. Toward that end, we first examined TEAD chromatin binding of across the genome and uncovered a striking redistribution of TEAD peaks from PRAD-specific to NEPC-specific enhancers, consistent with recent evidence that TEAD1 regulates gene expression in a human NEPC line (*48*). Equally striking was a parallel redistribution of FOXA1 peaks, raising the intriguing possibility that FOXA1, a pioneer factor expressed in both PRAD and NEPC, plays a pivotal role in initiating the lineage transition. Indeed, CRISPR deletion studies establish that FOXA1 is essential for the reshaping of the open chromatin landscape triggered by TEAD + NOTCH inhibition in PRAD organoids and enables the redistribution of TEAD peaks (see model, **Fig. 6F**).

Despite this new clarity, several questions remain regarding the mechanism by which RB functions as a gatekeeper of NE lineage plasticity. The fact that induction of the NE program can also occur in RB wildtype cells provides an important clue in that this induction is accompanied by activation of RB-dependent cell cycle arrest. Therefore, RB loss is required to overcome this cell cycle checkpoint to progress to NEPC. That said, a recent ChIP-seq study (performed in retinal pigment epithelial (RPE1) cells) revealed RB binding at enhancers, distinct from canonical RB binding sites in the promoters of genes that regulate cell cycle progression, raising the possibility that loss of this enhancer role of RB may also contribute to lineage plasticity (*49*). Interestingly, RB binding sites at these enhancers are enriched for TEAD motifs, providing a tantalizing potential connection to the ECM-integrin-YAP/TEAD axis defined here. The fact the efficiency of NE program induction differs across genotypes (and enhanced by TP53 and RB1 loss) is also of interest, reminiscent of similar genotype-specific differences in induced pluripotent stem (iPS) cell induction (*50*).

The fact that lineage plasticity in our models can be reversed by restoring YAP/TEAD activity through LATS kinase inhibition (or genetically through constitutive YAP activation) raises the intriguing possibility that CRPC patients at risk for plasticity could benefit from treatment with a clinical grade LATS inhibitor. Such an intervention might not only reinforce PRAD lineage maintenance but also reprogram tumors with mixed lineage phenotypes (an increasingly common clinical phenotype) to a homogeneous PRAD state. Because sustained systemic LATS inhibition may be poorly tolerated (based on hyperproliferation syndromes seen in LATS deficient mice) (*51*), evaluation of this “lineage restoration” approach in CRPC using LATSi would likely require transient therapy followed by treatment with definitive PRAD therapy. Recent clinical data in CRPC patients showing enhanced response to ARPIs through PRC2 complex inhibition is associated with restoration of luminal lineage gene expression, raising the possibility of alternative lineage restoration drug targets. Essential to the clinical success of any lineage restoration approach is the availability of an effective follow-on strategy to target the “restored” PRAD lineage state. Novel ARPIs might be one option (e.g., AR degraders), but emerging treatments using radioligands, antibody drug conjugates or T cell engagers directed at PRAD-specific cell surface antigens (such as PSMA or KLK2) may be more promising. It will also be important to explore whether the importance of YAP/TEAD activation in PRAD lineage maintenance reported here extends to other tumor settings in which lineage plasticity is also a mechanism of acquired resistance, such as EGFR-mutant lung cancer.

## Acknowledgements

We thank members of the Sawyers lab for valuable discussions and critical feedback. We are grateful to Lukas Dow and Cynthia Jung for advice and comments on preparation of the manuscript. We also acknowledge the staff of the MSKCC core facilities for their technical support, including the Molecular Cytology Core, Antitumor Assessment Core, Center for Epigenetics Research, Integrated Genomics Operation Core, and the Flow Cytometry Core. AJH was an Investigator of Howard Hughes Medical institute. Diagnosed in 2018 with high-grade prostate cancer, he underwent prompt, robot-assisted surgery at New York–Presbyterian Hospital, then received courteous, professional, and effective radiation treatment at Memorial Sloan-Kettering Cancer Center. While expressing his great gratitude to the personnel of both institutions, he approached prostate cancer in a spirit not of getting mad, but of getting even.

## Funding

TH is supported by the Prostate Cancer Research Program Early Investigator Research Award (W81XWH-21-PCRP-EIRA) from the Department of Defense and the Translational Research in Oncology Training (TROT) Program at MSKCC. ZS is supported by the Edith C. Blum Foundation Postdoctoral Training Award. PM is supported by ASCO Young Investigator Award and the Weinberg Fellowship. MCH is supported by P01CA298991 and P50CA097186. Integrated Genomics Operation (IGO) Core facility is funded by the NCI Cancer Center Support Grant (CCSG, P30 CA08748), Cycle for Survival, and the Marie-Josée and Henry R. Kravis Center for Molecular Oncology. CLS is supported by the Howard Hughes Medical Institute, Calico Life Sciences and NIH grants CA193837, CA092629, CA265768, CA008748.

## Author contributions

TH conceived the project, performed research, analyzed data and wrote the paper. ZS, ML and YC performed research and analyzed data. PM and MB performed research, analyzed data and wrote the paper. NK, SN, HZ, ES, SO, LF, WK, FN, FM, RP, JZ, NS, HK, NM, QC, ER and EC performed research. NS, PH, ED, ZC, MCH, DP, RK, and AJH contributed new reagents/analytic tools. CLS conceived the project, supervised the project and wrote the paper.

## Competing interests

CLS serves on the Board of Directors of BeOne and the scientific advisory boards of CellCarta, Column Group, Foghorn, Housey Pharma, Juri, Manas, Nextech, Nilo, PMV Pharma and ORIC.

## Data and materials availability

Sequencing data have been deposited to the Gene Expression Omnibus (GEO) database under accession number: GSE343498, GSE343500, GSE343502 and GSE343504. All other data is available in the manuscript or the supplementary materials.

## Supplementary Materials

## Materials and Methods

### Ethical statement

Mouse experiments were conducted under protocol 0607012 approved by the Institutional Animal Care and Use Committee of Memorial Sloan Kettering Cancer Center (MSKCC), New York.

### Organoid culture, analysis and transplantations

#### Mouse prostate organoid derivation and culture

Whole mouse prostates, including all lobes, were isolated as previously described (*52*, *53*). Briefly, dissected prostates were digested with collagenase type II (Gibco) for 2 hours at 37 °C, followed by TrypLE™ Express (Gibco) digestion at 37 °C until a single-cell suspension was obtained. All digestions were supplemented with Y-27632 (10 µM; MedChemExpress, HY-10583) to prevent anoikis, and the resulting cell suspension was filtered through 40-µm strainers to remove debris. Murine prostate organoids were established and maintained under standard culture conditions as previously described (*52*, *53*). In brief, dissociated epithelial cells were embedded in 20-µL drops of growth factor–reduced Matrigel (Corning, 356231) and overlaid with mouse prostate organoid medium.

#### Organoid genetic engineering

Genetic perturbation in organoids was performed using CRISPR–Cas9 ribonucleoprotein (RNP) complexes as previously described (*54*). Briefly, Cas9 protein (IDT) was incubated with synthetic sgRNA (IDT) to assemble the RNP complex prior to nucleofection. The sgRNA sequences are listed in table S1. A total of 1.2 µM cRNP was used per sgRNA. Dissociated organoid cells (5 × 10⁵–1 × 10⁶) were resuspended in nucleofection buffer containing RNP complexes and electroporation enhancer (IDT; 1:1 molar ratio to RNP) in a final volume of 100 µL. The suspension was transferred to a nucleofection cuvette and electroporated using a Lonza Amaxa Nucleofector II (program T-030). Cells were then centrifuged and seeded into Matrigel for culture. Pten-null organoids were enriched and maintained by culturing them in medium lacking EGF.

For lentiviral transduction, viral titers were determined prior to infection, and a multiplicity of infection (MOI) of 0.2 was used to ensure single-copy integration. A total of 5 × 10⁵ cells were used per reaction. cMyc overexpression was achieved by lentiviral transduction of a construct containing SFFV-cMyc*–*IRES*–*eGFP; GFP-positive cells were isolated by fluorescence-activated cell sorting (FACS).

For adenoviral transduction, 1 µL of Ad-CMV-Null (Vector Biolabs, 1300) or 1 µL of Ad-CMV-iCre (Vector Biolabs, 1045) was used to infect 5 × 10⁴ cells. To enhance infection efficiency, spinoculation was performed at 32 °C and 600 × *g* for 1 hour.

#### Ascl1 reporter organoids

*Ascl1*-CreER mice (JAX: 012882) were crossed with CAG-LSL-zsGreen reporter mice (JAX: 007906) to generate *Ascl1*-CreER; CAG-LSL-zsGreen mice. Prostate organoids were derived from these mice, and sequential tumor suppressor knockouts were generated using RNP-mediated CRISPR editing as described above. Tumor suppressor loss was validated by Western blotting. To induce CreER-mediated recombination and zsGreen expression in *Ascl1*-expressing cells, organoids were treated with 1 μM 4-hydroxytamoxifen (4-OHT; MedChemExpress, HY-16950) for 24 h.

#### RNA isolation, cDNA Synthesis and quantitative PCR

RNA was isolated from organoids using the RNeasy Plus Mini Kit (Qiagen) according to manufacturer’s protocol. RNA concentration was quantified using a NanoDrop (ThermoFisher). Complementary DNA (cDNA) was synthesized using High-Capacity cDNA Reverse Transcription Kit (Thermo Scientific) according to manufacturer’s instructions. Quantitative PCR experiments were conducted on Applied Biosystems QuantStudio 6 Flex Real-Time PCR system. To capture the transcript with low abundance, we performed qPCR using 50 amplification cycles rather than the standard 40. Melting curve analysis was routinely used to confirm specificity of the amplified products. qPCR primers were listed in table S1.

#### Suspension culture

Organoid cells were dissociated using TrypLE^TM^ Express Enzyme (Gibco) and passed through 40um cell strainers (Fisher Scientific). A total of 1 x 10^6^ cells were resuspended in prostate organoid medium and seeded into one well of a Nunclon^TM^ Sphera^TM^ plate (Thermo Scientific, 174932) or an Ultra-low attachment plate (Corning, 3471). Cells were passaged every 3-4 days. For passaging, cells were enzymatically dissociated to single cells using TrypLE^TM^ Express prior to reseeding.

#### Protein isolation and western blot analysis

Organoids were isolated from the Matrigel using cell recovery solution (Corning, 354253). Cells were lysed in RIPA buffer containing protease inhibitors (Calbiochem) and phosphatase inhibitors (Calbiochem). Protein concentrations were quantified using a bicinchoninic acid (BCA) assay (Pierce, Thermo Fisher). Lysates were denatured using 4X protein loading dye (SDS 200 nM Tris, 8% SDS, 0.4% bromophenol blue, 40% glycerol, 400 mM 2-mercaptoethanol, pH 6.8). 20-30 µg of protein was loaded on NuPage 4-12% gradient polyacrylamide gels (Invitrogen). After electrophoresis, protein was transferred to a PVDF membrane and blocked with 5% milk in TBS-T. Primary antibodies were incubated overnight at 4 degrees. Membranes were washed using TBS-T and incubated with secondary antibodies for 1 h at room temperature with shaking. Proteins were visualized using ECL prime (Amersham, GE healthcare) and ImageQuant 800 (Amersham, GE healthcare). Antibodies used in this study are listed in the table S1.

#### Orthotopic transplantation

Organoid cells (2 x 10^5^) were resuspended in 20 μl of a 1:1 mixture of growth factor-reduced Matrigel (Corning, 356231) and organoid culture medium and injected into the prostate dorsal lobes of immunodeficient NSG mice (JAX 005557) or C57BL/6J mice (JAX 000664) at 2 months of age.

#### Subcutaneous transplantation

Organoid cells (2 x 10^5^) were resuspended in 100 μl of a 1:1 mixture of growth factor-reduced Matrigel (Corning, 356231) and organoid culture medium and injected subcutaneously into the right flank of immunodeficient NSG mice (JAX 005557) or C57BL/6J mice (JAX 000664) at 2 months of age.

#### Generation of NEPC tumoroids

Mouse prostate tumors were dissected and finely minced using sterile surgical blades. Tumor fragments were enzymatically dissociated in collagenase type II (Gibco) for 1 hour at 37 °C, followed by further digestion with TrypLE™ Express (Gibco) for 30 minutes at 37 °C. All digestions were supplemented with Y-27632 (10 µM) to prevent anoikis. The resulting cell suspension was passed through 40-µm cell strainers (Fisher Scientific) to obtain a single-cell suspension. Cells were stained on ice for 1 hour with CD51-PE (1:200; BioLegend, 104106) and EpCAM-Alexa Fluor 647 (1:200; Abcam, ab237385). CD51⁻/EpCAM⁺ cells were isolated by fluorescence-activated cell sorting (FACS) and cultured under standard prostate organoid culture conditions.

### Histology and immunostaining

For GEMM experiments, whole mouse prostates containing all lobes were collected at autopsy. Histological data from all lobes were pooled for quantification. For intraprostatic transplantation and adenoviral infection experiments, injected prostate lobes were collected at autopsy. Prostate tissues were fixed using 4% paraformaldehyde, dehydrated with 70% ethanol, paraffin-embedded and sectioned. H&E staining was performed following standard protocols by the MSKCC Molecular Cytology Core. Immunohistochemistry and immunofluorescence were performed on a Leica Bond RX automatic stainer using antibodies listed in the table S1. Formalin-fixed, paraffin-embedded (FFPE) stained tissue sections were scanned using a Pannoramic P250 Flash scanner (3DHISTECH, Hungary) equipped with a 20×/0.8 NA objective. Whole-slide images were exported as .tif files using *SlideViewer* software (3DHISTECH, Hungary) for subsequent analysis in *ImageJ/FIJI* (NIH, USA). For immunohistochemistry (IHC) quantification, color deconvolution was applied to separate hematoxylin and DAB signals. The tissue area and DAB-positive area were determined by thresholding, and the ratio of DAB-positive area to total tissue area was used as a quantitative measure of staining intensity. For immunofluorescence (IF) quantification, multi-channel images were similarly exported and analyzed in *ImageJ/FIJI*. Tissue area was defined using the DAPI channel. Nuclei were segmented by generating a DAPI mask followed by watershed separation. Regions of interest (ROIs) approximating individual cells were then used to assess marker positivity, defined by the percent area of signal per marker channel relative to the nuclear ROI.

### In situ immunolabeling of YAP1 and ASCL1 in human prostate cancer tissues

Formalin-fixed, paraffin-embedded (FFPE) metastatic prostate cancer specimens were procured through the University of Washington Rapid Autopsy Program as previously described (*55*, *56*). For sequential ASCL1/YAP1 co-immunofluorescence, FFPE tissue sections were deparaffinized and subjected to antigen retrieval by steaming in 10 mmol/L sodium citrate buffer (pH 6.0; Vector Laboratories) for 30 minutes. Slides were first incubated with ASCL1 primary antibody (Abcam, ab211327; 1:50 dilution), followed by detection using Tyramide-Alexa Fluor 488 (Life Technologies, B40953). Antibody stripping was subsequently performed by steaming slides for 15 minutes in Target Retrieval Solution (pH 9.0; Dako). Slides were then incubated with YAP1 primary antibody (Abcam, ab205270; 1:300 dilution), followed by anti-rabbit IgG secondary antibody detection (Leica Microsystems, PV6119) and visualization using Tyramide-Alexa Fluor 568 (Life Technologies, B40956). Nuclear counterstaining was performed using DAPI.

### Bulk RNA-seq

RNA was extracted using the RNeasy Plus Mini Kit (Qiagen) from bulk samples and sequenced at the Integrated Genomics Operation Core (MSKCC). cDNA was synthesized from purified RNA using oligo(dT) primers and reverse transcriptase according to standard Illumina protocols. The resulting cDNA was subjected to automated Illumina paired-end library construction. Libraries were sequenced on Illumina HiSeq2000 instruments with paired reads of 100 base pairs (bp) in length per sample, generating approximately 30–40 million reads per sample. Sequence data were processed and analyzed using Partek^TM^ Flow^TM^ software, v11.0. Briefly, raw FASTQ files were quality-checked and trimmed to remove low-quality bases and adaptor sequences. Cleaned reads were aligned to the mouse reference genome (mm10) using the STAR aligner with default parameters. Aligned reads were quantified at the gene level based on Ensembl transcript annotations. Gene-level read counts were normalized using the Fragments Per Kilobase of transcript per Million mapped reads (FPKM) method to account for sequencing depth and transcript length. For visualization, normalized expression values were log2-transformed. Heatmaps and principal component analysis (PCA) plots were generated using normalized expression values to visualize global transcriptomic differences among samples. Differential gene expression was analyzed using the DESeq2 algorithm (v1.34.0). Genes with an adjusted *p* < 0.05 and absolute log2 fold change > 2 were considered significantly differentially expressed. Gene set enrichment analysis (GSEA) was performed using the GSEA software (https://www.gsea-msigdb.org/gsea/index.jsp). The PRAD and NEPC gene sets used in this study are listed in table S2.

#### PRAD-to-NEPC transition score analysis

Raw counts from bulk RNA-seq were normalized to counts per million (CPM) to account for differences in sequencing depth between samples, followed by log2 transformation after adding a pseudo count of one. Genes were retained if they reached at least 1 CPM in at least three samples. The adenocarcinoma reference state was defined as the average expression profile of the three DMSO samples. The neuroendocrine reference state was defined as the average expression profile of all nine NE samples, including three biological replicates each from the NEPC-1, NEPC-2, and NEPC-3. Using all three NE models to define the neuroendocrine reference reduces the influence of model-specific transcriptional features and captures the transcriptional program shared across independent NEPC models. The difference between the adenocarcinoma and neuroendocrine reference expression profiles defines a transcriptional axis representing the direction of lineage transition. Each sample was projected onto this axis to determine its position relative to the two reference states. The projection was scaled such that the adenocarcinoma reference centroid had a score of 0 and the neuroendocrine reference centroid had a score of 1.

### Bulk RNA-seq analysis on patient cohort

RNA-seq data from two clinical cohorts were obtained from the cBioPortal for Cancer Genomics (*57*). The cohorts included: the SU2C/PCF Dream Team cohort (210 adenocarcinoma and 22 neuroendocrine prostate cancer samples) (*12*), and the Beltran *et al.* cohort (*23*).

YAP/TAZ target score and Integrin score were calculated using Gene Set Variation Analysis (GSVA) with the single-sample Gene Set Enrichment Analysis (ssGSEA) method (*58*), implemented in R (version 4.3.2). The YAP/TAZ target gene set was derived from Wang et al (*59*), and the Integrin score was calculated using all annotated integrin genes as the input gene set. Statistical differences in gene set scores and mRNA expression levels of individual genes between adenocarcinoma and NEPC samples were assessed using a two-tailed Wilcoxon rank-sum test.

### Visium spatial transcriptomics

#### Spatial transcriptomics acquisition and preprocessing

Visium spatial transcriptomics data were downloaded from GEO accession GSE278936, including spatial alignments, high-resolution histology images, and spot-level gene expression matrices. After loading the data into Scanpy, low-quality spots (<500 total UMIs) and genes expressed in fewer than three spots were filtered on a per-sample basis. Gene counts were deduplicated by summing counts for features sharing the same gene symbol.

#### Single-cell reference atlas

Single-cell RNA sequencing data comprising 119,083 cells from prostate tumors were obtained from Zaidi et al. (2024) (*29*) (“Single-cell analysis of treatment-resistant prostate cancer”) as downloaded from GEO (GSE264573). This dataset includes annotations for major cell types, including myeloid, lymphoid, stromal, normal epithelial, and tumor epithelial cells. Tumor epithelial cells in this dataset are further subclassified into castration-sensitive prostate cancer (CSPC), castration-resistant prostate cancer (CRPC), and neuroendocrine prostate cancer (NEPC).

#### Cell type deconvolution using BayesPrism

Because Visium spots capture transcripts from multiple cells, spot-level deconvolution was performed to infer underlying cell type composition in the Kiviaho et al. (2024) (18) spatial dataset. Reference-based spatial deconvolution was performed using BayesPrism.

To improve deconvolution accuracy, a feature selection step was performed following recommendations from the BayesPrism authors. Marker genes were identified from the Zaidi et al. (2024) (*29*) single-cell reference by performing pairwise differential gene expression analysis between cell types using Scanpy’s rank_genes_groups function with the “t-test_overestim_var” method applied to log-normalized gene expression values. Genes were retained if they exhibited a log fold change ≥ 0.25 and an adjusted p-value < 0.05 across all relevant pairwise comparisons.

This feature selection strategy was chosen to reduce noise from non-informative genes and improve deconvolution stability.

The number of retained marker genes per cell type or state was as follows: CSPC (1,846), NEPC (7,786), CRPC (3,314), non-tumor epithelial (1,336), lymphoid (1,405), myeloid (2,195), and stromal (2,334).

Spot-level raw gene counts from each Visium sample and cell-level raw counts from the single-cell reference atlas were used as inputs to BayesPrism (pybayesprism). For each sample, expression matrices were restricted to genes shared between the spatial data and the selected marker gene lists.

BayesPrism yields relative transcriptional contribution (“theta”) estimates for each cell state per spot, which were used in all downstream analyses.

#### BayesPrism deconvolution robustness analysis via downsampling

To assess the robustness of BayesPrism deconvolution given the sparsity of Visium data, we performed UMI downsampling analyses. For each of the 48 Visium slides, BayesPrism deconvolution was repeated 20 times following random downsampling of 10% of UMIs per slide, retaining 90% of UMIs. Pairwise Spearman correlations (190 comparisons per sample) were computed between theta estimates derived from downsampled runs for each cell type. Spearman correlation was chosen due to the non-normal distribution of theta estimates.

#### Pathologist annotation–based spot-level validation

To validate deconvolution results, spot-level pathological annotations provided by the authors of Kiviaho et al. (2024) (18) were compared with BayesPrism estimates. Pathologist annotation categories included Stroma, Benign, Inflammation, Atrophy, PIN, and tumor-associated spots annotated by Gleason score (e.g., Gleason X). Notably, prostate tumor subclasses such as NEPC or neuroendocrine-like pathology were not included in the pathological annotations and therefore could not be independently assessed.

Of 112,080 spots retained following quality control and filtering, 25,186 spots had corresponding pathologist annotations. For these spots, the total deconvolved tumor fraction (defined as the sum of CSPC, CRPC, and NEPC theta estimates) and the deconvolved fibroblast fraction were compared with pathological annotations.

#### Analysis of spatial co-localization of tumor and stroma

A spot was classified as NEPC-high, CRPC-high, CSPC-high, or stroma-high if its corresponding theta estimate exceeded 0.25. PRAD-high spots were defined by a combined CRPC+CSPC theta >0.25. Spots exceeding this threshold for both a tumor cell state and stromal content were classified as mixed. To assess sensitivity to threshold choice, tumor theta thresholds were varied from 0.1 to 0.4, and downstream analyses were repeated.

#### Stratification by tissue origin

To address the potential confound of comparing primary and metastatic tissue, spatial co-localization analyses were also performed in metastatic sites of disease separately from localized.

The fraction of NEPC-high and PRAD-high spots co-localizing with high stromal content was computed and compared using Fisher’s exact test.

### Bulk RNA-seq deconvolution

To assess stromal content across a larger cohort of advanced prostate cancer specimens, the same BayesPrism deconvolution framework was applied to bulk RNA-seq data from the SU2C/PCF cohort (Abida et al. 2019, n=266). Poly-A selected FPKM expression data and clinical annotations (including neuroendocrine features, tissue site, and pathology classification) were obtained from cBioPortal (prad_su2c_2019). Expression matrices were restricted to genes shared between the bulk data and the marker gene set (11,813 genes retained). BayesPrism was run with identical parameters as the spatial analysis, yielding per-sample cell state fractions. BayesPrism deconvolution results were validated against the published neuroendocrine annotation (NEUROENDOCRINE_FEATURES), confirming significantly higher NEPC fractions in NE-positive samples (mean theta 0.350 vs 0.052, Mann–Whitney P = 1.14 × 10⁻⁶).

### Stromal fraction comparison by metastatic site

SU2C samples were classified as NEPC or CRPC-adenocarcinoma using the published pathology classification (Small cell and Adenocarcinoma with NE features vs Adenocarcinoma; samples with inadequate or unavailable pathology were excluded). Stromal fractions (BayesPrism theta) were compared between groups using the Mann-Whitney U test. Annotated metastatic site was also used to further stratify the pattern of NEPC localization and derive a median stromal fraction per site.

### Single-cell multiome

#### Data pre-processing

The FASTQ files of the single-cell multiome data were processed by sample with cellranger-arc v. 2.0.2 with alignment against reference genome mm10. All samples were aggregated using the cellranger-arc aggr function v. 2.0.2. The resulting consensus peak list of genomic regions was annotated with the HOMER (*60*) function version 4.11 with the cellranger-arc refdata-cellranger-arc-mm10-2020-A-2.0.0 genes.gtf file.

#### Quality control

Cell filtering criteria were calculated independently for each sample and modality. For single-cell gene expression data, we determined the library size per cell, the number of genes expressed per cell, and the fraction of mitochondrial counts per cell. Specifically, cells expressing fewer than 1,000 or more than 10,000 genes, those with fewer than 1,000 or more than 20,000 total UMI counts, or those exhibiting a mitochondrial read fraction exceeding 20% were excluded. Following this initial cellular filtration, a doublet detection score was computed using Scrublet (*61*) v. 0.2.3, employing the parameters min_counts = 2, min_cells = 3, vscore_percentile = 85, n_pc = 50, expected_doublet_rate = 0.02, sim_doublet_ratio = 3, and n_neighbors = 15. Subsequently, all cells with a doublet score greater than 0.14 were removed. Finally, genes expressed in fewer than 3 cells were discarded.

For single-cell chromatin accessibility data, we assessed the minimum library size per cell, the number of peaks detected per cell, the maximum TSS enrichment score, and the nucleosome signal (defined as the ratio of nucleosome-free to mono-nucleosome fragments per cell). Specifically, cells with fewer than 500 or more than 20,000 peaks, or with a total UMI count below 1,000 or above 40,000, were excluded. Additionally, all peaks detected in fewer than 50 cells were removed. Cells were filtered separately for gene expression and chromatin accessibility, with only those cells passing quality control in both modalities being retained. The resulting dataset comprised 23,134 cells, 23,312 genes, and 147,895 peaks.

#### Normalization and feature selection

For single-cell GEX, normalization was performed using a shifted log-transformation of counts, divided by the library size and scaled to 10,000 reads (logCPM+1). For single-cell ATAC normalization, Term Frequency - Inverse Document Frequency (TF-IDF) was computed to weigh the importance of open chromatin regions. Specifically, a shifted log-transformation of the term frequency (TF) was utilized, a method demonstrated to be beneficial in sparse datasets compared to direct TF application, as implemented in the muon (*62*) ATAC module.

To select highly variable genes, the scanpy (*63*) (v. 1.10.3) function pp.compute_highly_variable was employed with the ‘cellranger’ flavor, resulting in the selection of 3,000 highly variable genes per sample. Highly variable peaks were computed using the scanpy function pp.compute_highly_variable, with parameters min_mean=0.05 and min_disp=0.5, yielding 16,865 highly variable peaks.

Cell cycle genes were scored to determine the cell cycle phase using the scanpy tl.score_genes_cell_cycle function per sample. As a reference, the human cell cycle gene list curated by Tirosh et al. (*64*) was converted to align with the mouse gene symbol convention.

#### Dimensionality reduction

For dimensionality reduction, principal component analysis (PCA) was applied to gene expression data, and latent semantic indexing (LSI) (*65*) was utilized for chromatin accessibility. A weighted nearest neighbor (WNN) graph, incorporating L2-regularization of distances derived from both modalities on highly variable features, was then computed to generate a joint UMAP.

#### Batch correction

We did not perform any batch correction in the final dataset, because all samples were processed as a single batch.

#### Clustering and annotation

Leiden (*66*) clustering, at a resolution of 0.5, resulted in seven distinct clusters. Gene characterization was subsequently performed utilizing a t-test within the scanpy tl.rank_genes_groups function, employing a one-versus-rest comparison. Differentially accessible peaks were characterized using the Wilcoxon rank-sum test, also within the scanpy tl.rank_genes_groups function and a cluster-versus-rest setup. The top 20 genes/peaks were then investigated to characterize each cluster. The smallest cluster, comprising 444 cells from all three samples, was subsequently excluded.

All analyses were performed in Python v. 3.10.15.

### Chromatin profiling and data analysis

#### Bulk ATAC-seq

Freshly harvested prostate organoid cells were processed by MSKCC’s Epigenetics Research Innovation Lab. ATAC was performed using 50,000 cells per replicate as previously described (*67*) using the OpenTn5 enzyme (*68*). The sequencing libraries were purified with SPRIselect magnetic beads (B23318, Beckman Coulter), quantified using a Qubit Flex fluorometer (ThermoFisher Scientific) and profiled using a TapeStation 4200 (Agilent). The libraries were sequenced by the MSKCC Integrated Genomics Operation core facility. After PicoGreen quantification and quality control by Agilent TapeStation, libraries were pooled and run on a NovaSeq 6000 in a PE100 run, using the NovaSeq 6000 S4 Reagent Kit (200 Cycles) (Illumina). The loading concentration was 0.5nM and a 1% spike-in of PhiX was added to the run and for quality control purposes. The run yielded on average 50-60M reads per sample.

#### CUT&RUN

Freshly harvested prostate organoid cells were processed by MSKCC’s Epigenetics Research Innovation Lab. CUT&RUN was performed with 1M cells per replicate using the CUTANA™ ChIC/CUT&RUN Kit (Epicypher #14-1048) and the following antibodies: rabbit polyclonal anti-FOXA1 (Invitrogen PA5-27157); rabbit monoclonal anti-YAP1 (CST, #14074); rabbit monoclonal anti-TEAD1 (CST, #12292); rabbit anti-H3K4me3 (Epicypher, #13-0041); rabbit anti-mouse IgG (Epicypher, #13-0042). The recovered DNA fragments were quantified and sent to the MSKCC Integrated Genomics Operation core facility for library preparation and sequencing. Immunoprecipitated DNA was quantified by PicoGreen and the size was evaluated by Agilent BioAnalyzer. Illumina sequencing libraries were prepared using the KAPA EvoPrep Kit (Roche 10212250702) according to the manufacturer’s instructions with 0.2-5 ng input DNA and 14 cycles of PCR. Barcoded libraries were run on the NovaSeq 6000 in a PE100 run, using the NovaSeq 6000 S4 Reagent Kit (200 Cycles) (Illumina). An average of 5-10M paired reads were generated per sample.

#### Sequencing data analysis

Raw sequencing reads were trimmed and filtered for quality (Q>15) and adapter content using version 0.4.5 of TrimGalore (https://www.bioinformatics.babraham.ac.uk/projects/trim_galore) and running version 1.15 of cutadapt and version 0.11.5 of FastQC. Version 2.3.4.1 of bowtie2 (http://bowtie-bio.sourceforge.net/bowtie2/index.shtml) was employed to align reads to mouse assembly mm10 and alignments were deduplicated using MarkDuplicates in Picard Tools v2.16.0. Enriched regions were discovered using MACS2 (https://github.com/taoliu/MACS) with a p-value setting of 0.001, filtered for blacklisted regions (http://mitra.stanford.edu/kundaje/akundaje/release/blacklists/mm10-mouse/mm10.blacklist.bed.gz), and a peak atlas was created using +/- 250 bp around peak summits for ATAC data or using the entire ‘narrowPeak’ region for CUT&RUN data. The BEDTools suite (http://bedtools.readthedocs.io) was used to create normalized bigwig files. Version 1.6.1 of featureCounts (http://subread.sourceforge.net) was used to build a raw counts matrix and DESeq2 was employed to calculate differential enrichment for all pairwise contrasts for samples with replicates. For single sample data, MACS2 was run by swapping bams of different conditions to find differential regions. Sample concordance was evaluated by calculating intra- vs -inter-group variance in principal component analysis. Clusters were discovered by creating a superset of all differential peaks then running k-means clustering from k=2:10 until cluster redundancy emerged. Peak-gene associations were created by assigning all intragenic peaks to that gene, while intergenic peaks were assigned using linear genomic distance to transcription start sites (TSS). Pathway enrichment was calculated by assigning each gene a unique score based on the associated peak with the greatest magnitude change and running GSEA in pre-ranked mode. Motif signatures were obtained using Homer v4.5 (http://homer.ucsd.edu) on differentially enriched peak regions. Composite and tornado plots were created using deepTools v3.3.0 by running computeMatrix and plotHeatmap on normalized bigwigs with average signal sampled in 25 bp windows and flanking region defined by the surrounding 2 kb. Network analysis was performed using enrichplot::cnetplot in R with default parameters. Integrative analysis was performed by creating a merged superset of ATAC and CUT&RUN peaks and re-computing differential peaks over the new atlas, followed by projecting differential expression using the peak-gene annotations.

---

**Supplemental Figure 1.**
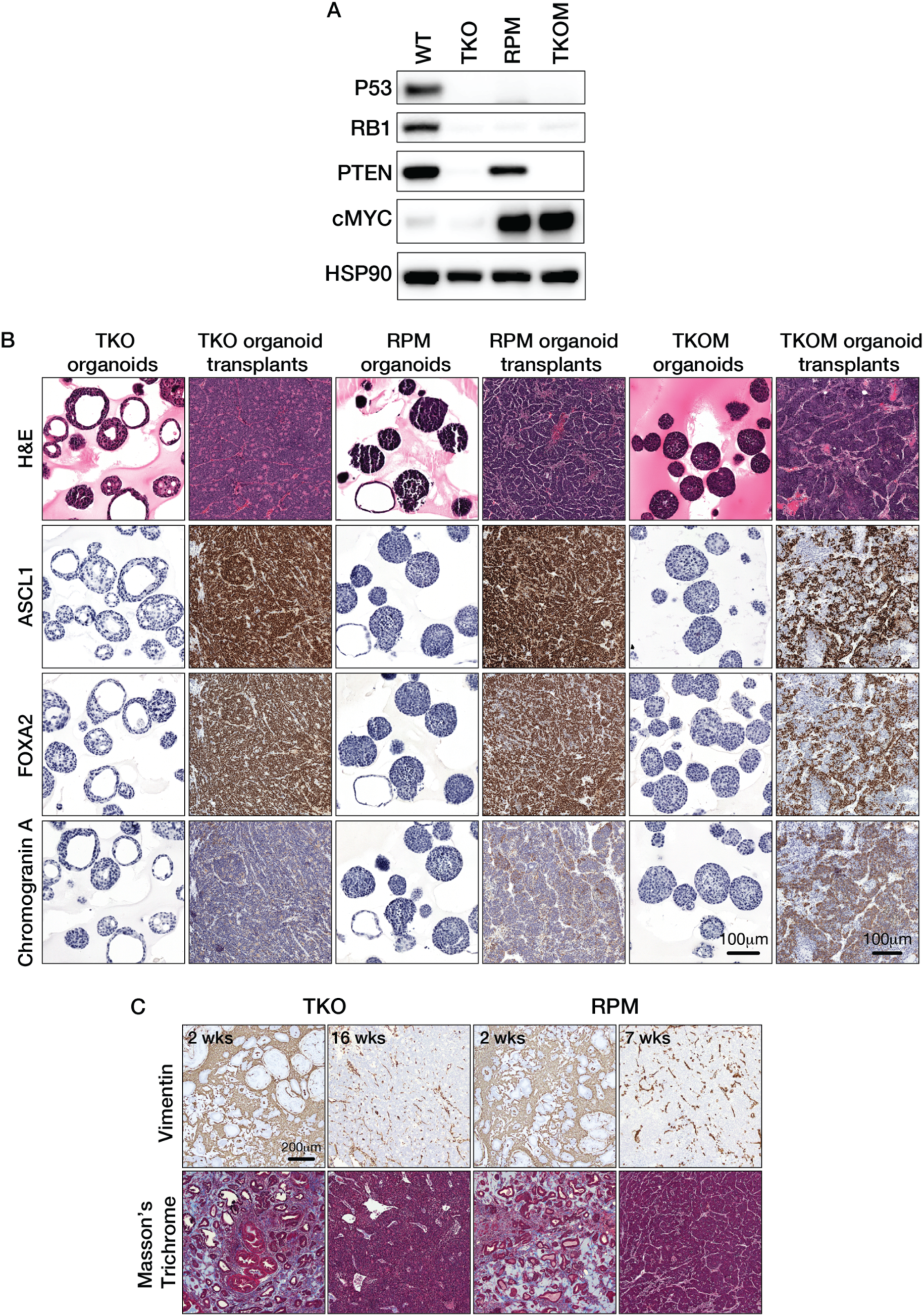
Cancer-associated fibroblasts and ECM are depleted in NEPC. (A) Western blot analysis confirming disruption of *Trp53*, *Rb1* and *Pten*, as well as overexpression of *cMYC* in engineered mutant organoid lines. (B) Hematoxylin and eosin (H&E) staining and immunohistochemistry (IHC) for ASCL1, FOXA2, and Chromogranin A in TKO, RPM, and TKOM organoids and their corresponding orthotopic tumors. (C) Vimentin IHC and Masson’s trichrome staining of early- and late-stage orthotopic transplants derived from TKO and RPM organoids.

**Supplemental Figure 2.**
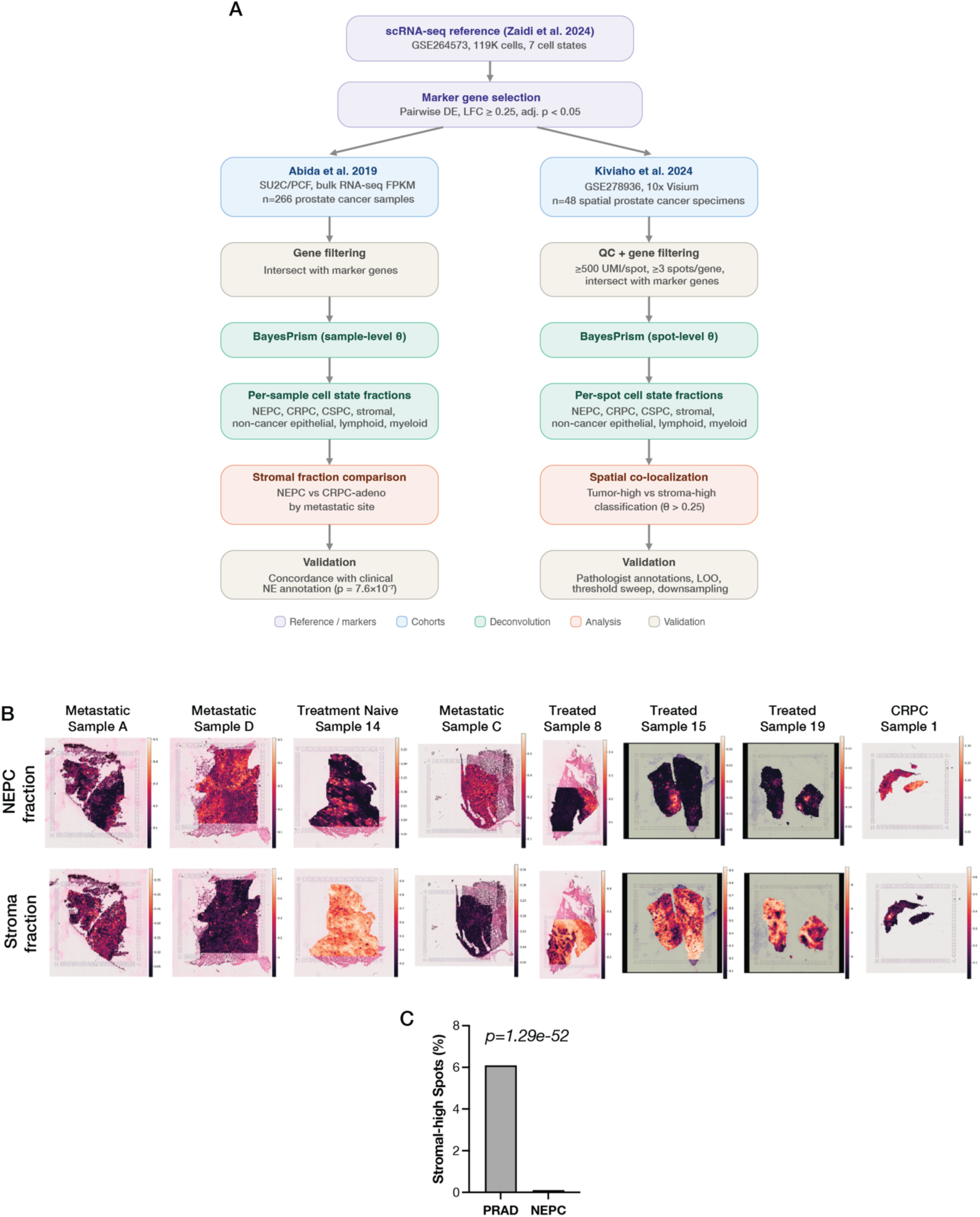
Tumor stroma is depleted in human NEPC. (A) Overview of the BayesPrism deconvolution and stromal analysis pipeline. A shared single-cell RNA-seq reference atlas (Zaidi et al. 2024; 119,083 cells, 7 cell states) was used to derive marker genes and perform deconvolution across two independent cohorts. (Left) Bulk analysis: RNA-seq from the SU2C/PCF cohort (Abida et al. 2019; n=266 samples) was deconvolved at the sample level, followed by stromal fraction comparison between NEPC and CRPC-adenocarcinoma stratified by metastatic site. Deconvolution was validated against published clinical neuroendocrine annotations (p = 7.6×10⁻⁷). (Right) Spatial analysis: Visium spatial transcriptomics data from Kiviaho et al. 2024 (n=48 specimens) were deconvolved at the spot level, followed by tumor-stroma co-localization analysis and validation against pathologist annotations, leave-one-out analysis, threshold sensitivity, and UMI down sampling. (B) Deconvolved NEPC theta estimates and corresponding stromal theta estimates across eight additional specimens (see also Fig 1D). Spatial maps highlight the distribution of NEPC and fibroblast-like content across Visium spots. (C) Quantification of stromal content in metastatic samples from the same cohort shown in (B).

**Supplemental Figure 3.**
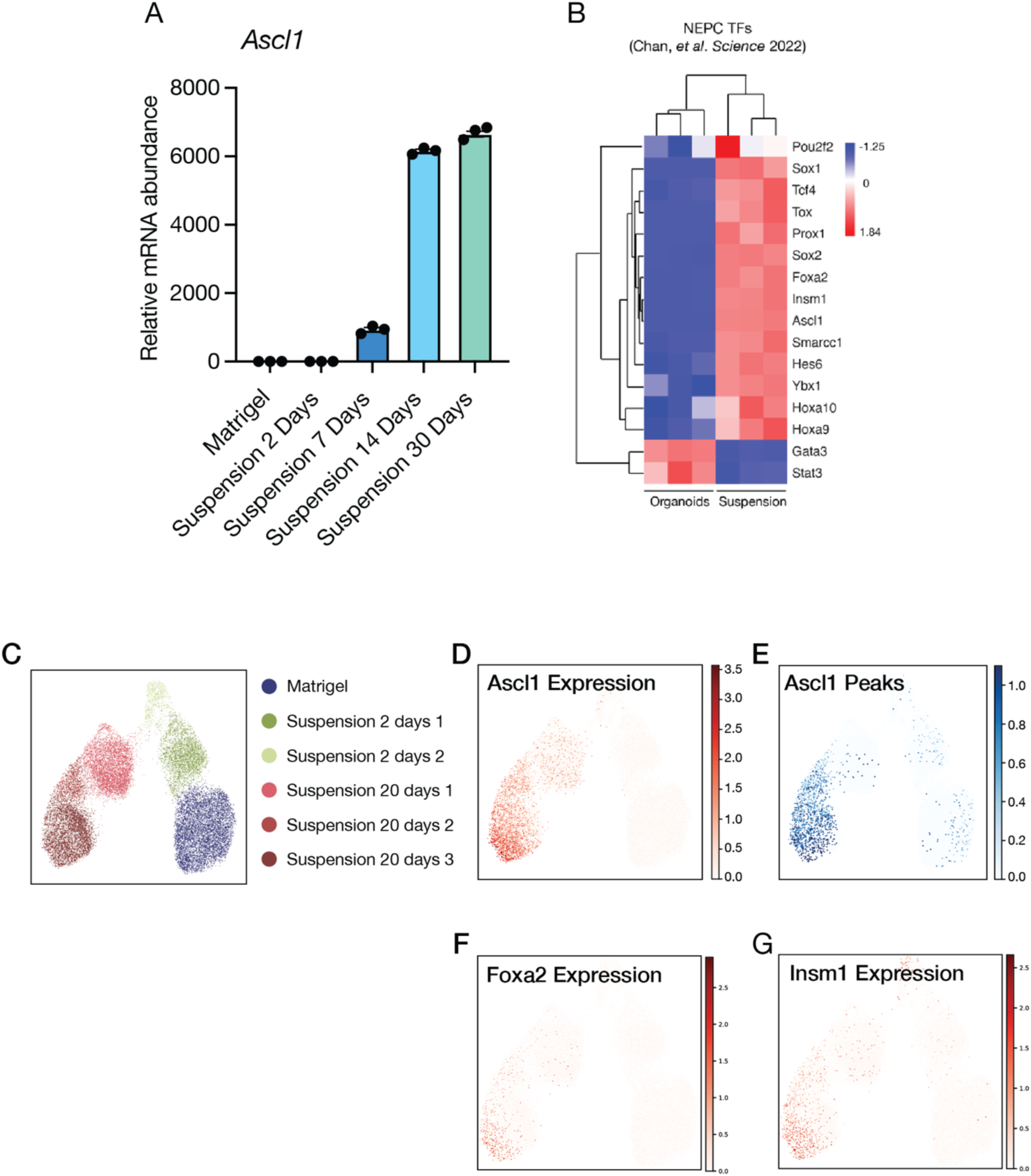
Removal of extracellular matrix induces Ascl1 expression. (A) Time-course qRT-PCR analysis showing progressive induction of Ascl1 in TKOM organoids cultured under suspension conditions. Error bars represent ±SEM, n=3. (B) Heatmap displaying expression levels of NEPC transcription factors in TKOM organoids cultured in Matrigel versus suspension. (C) UMAP of scRNA-seq data from TKOM organoids under the indicated conditions. (D-G) UMAPs showing Ascl1 expression (D), Ascl1 peaks (E), Foxa2 expression (F) and Insm1 expression (G) in TKOM organoids, corresponding to the conditions in (C).

**Supplemental Figure 4.**
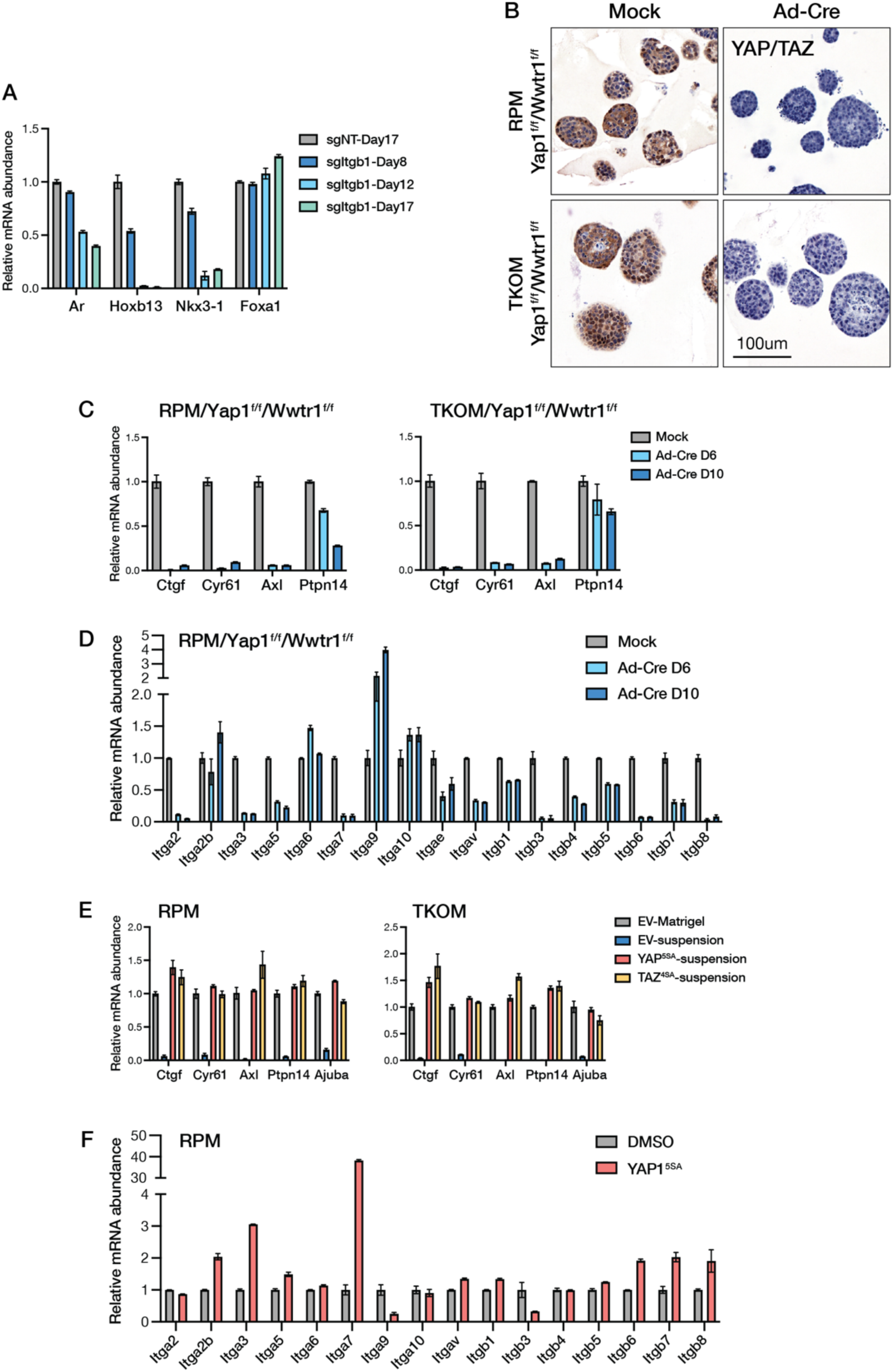
The integrin-YAP1/TEAD axis gatekeeps the PRAD state and restrains NEPC transition. (A) qRT-PCR analysis of PRAD lineage marker expression in sgNT and sgItgb1 organoids at the indicated time points following *Itgb1* knockout. Data are shown as mean ± SEM, n=3. (B) Representative YAP/TAZ IHC images of RPM and TKOM organoids harboring conditional *Yap1* and *Wwtr1* alleles, with or without Ad-Cre–mediated deletion. Scale bar, 100 μm. (C) qRT-PCR analysis of YAP1/TAZ target genes in RPM and TKOM organoids at 6 and 10 days following Yap1 and Wwtr1 deletion via Adeno-Cre (Ad-Cre) infection. Error bars represent ±SEM, n=3. (D) qRT-PCR analysis of integrin gene expression in RPM organoids at 6 and 10 days following Yap1 and Wwtr1 deletion via Ad-Cre. Error bars represent ±SEM, n=3. (E) qRT-PCR analysis of YAP1/TAZ target genes in RPM and TKOM organoids culturing in Matrigel, or in suspension for 14 days while overexpressing either empty vector (EV), constitutively active YAP (YAP^5SA^) or constitutively active TAZ (TAZ^4SA^). Error bars represent ±SEM, n=3. (F) qRT-PCR analysis of integrin gene expression in RPM organoids 24h after YAP^5SA^ overexpression. Error bars represent ±SEM, n=3.

**Supplemental Figure 5.**
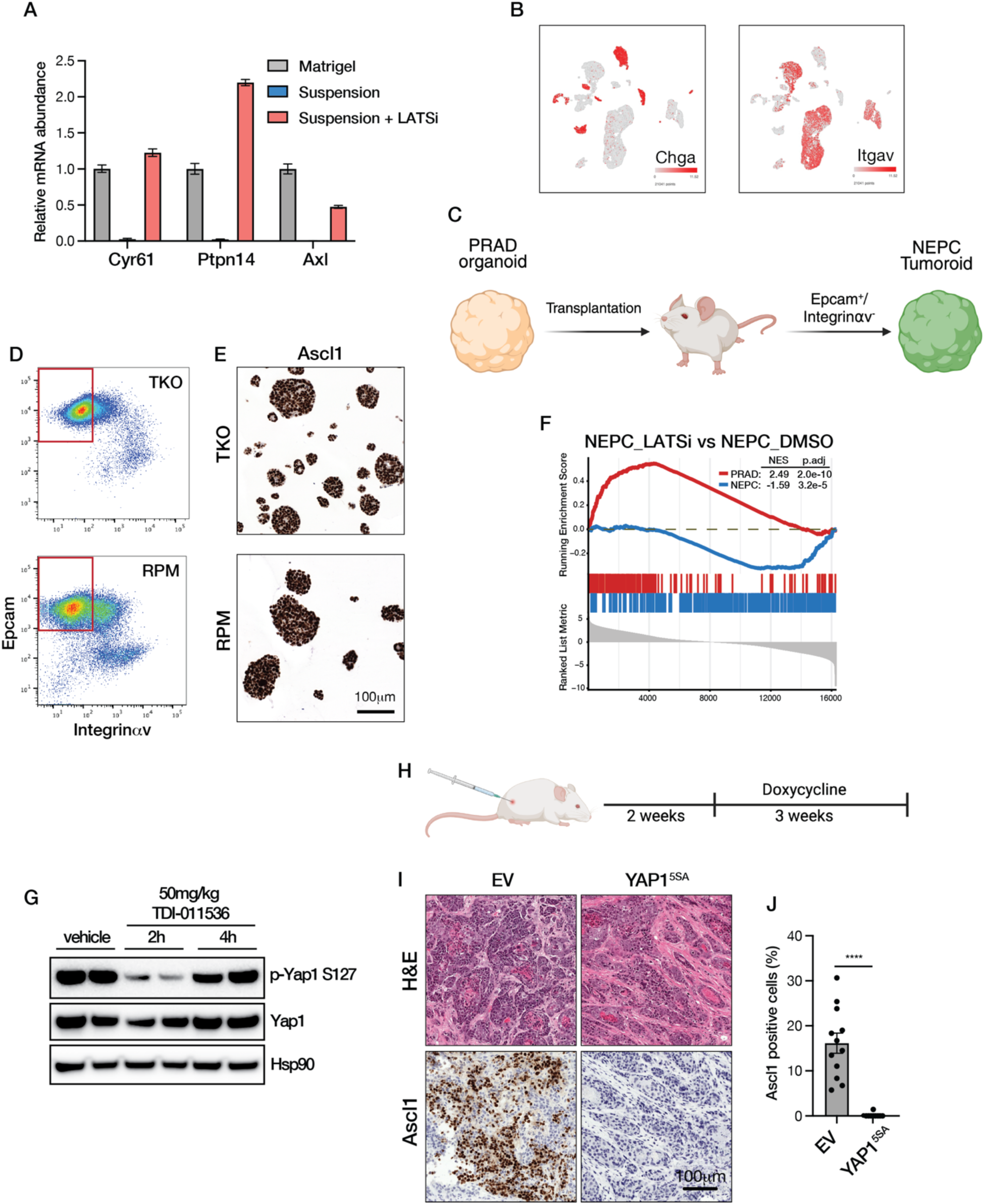
LATS inhibition impairs acquisition and maintenance of neuroendocrine state. (A) qRT-PCR of YAP/TAZ target genes (*Cyr61, Ptpn14 and Axl*) in TKOM organoids cultured in Matrigel, suspension, or suspension plus LATSi (5uM TRULI) for 14 days. Error bars represent ±SEM, n=3. (B) UMAP of epithelial cells in TKO GEMM tumors by scRNA-seq, showing that *Itgav* is silenced in NEPC cells but highly expressed in adenocarcinoma cells. NEPC cells are positive for *Chga*, whereas adenocarcinoma cells are *Chga*-negative. (C) Schematic of the workflow used to generate NEPC tumoroids from PRAD organoids. (D) Representative FACS plots showing sorting of Epcam^+^/Integrinαv^-^ cells from TKO or RPM tumors. (E) Representative Ascl1 IHC images of organoids derived by sorting Epcam^+^/Integrinav^-^ cells from TKO or RPM orthotopic tumors. The scale bar represents 100um. (F) Gene set enrichment analysis of ATAC-seq data comparing LATSi- and DMSO-treated NEPC organoids, showing enrichment of PRAD- and NEPC-associated gene sets. Normalized enrichment scores (NES) and adjusted *P* values are indicated. The PRAD and NEPC gene sets are listed in table S2. (G) Western blot analysis of phospho-Yap1 (S127) in lysates from mouse prostates following treatment with vehicle or TDI-011536 for 2 or 4 hours. (H) Schematic showing the timeline of the subcutaneous transplantation experiment using RPM organoids. (I) Representative H&E images and Ascl1 IHC images of RPM transplantation tumors overexpressing EV or YAP1^5SA^, n= 12 for each group. (J) Quantification of the percentage of Ascl1 positive cells in (I). Error bars represent ±SEM, n=12, ****p<0.0001.

**Supplemental Figure 6.**
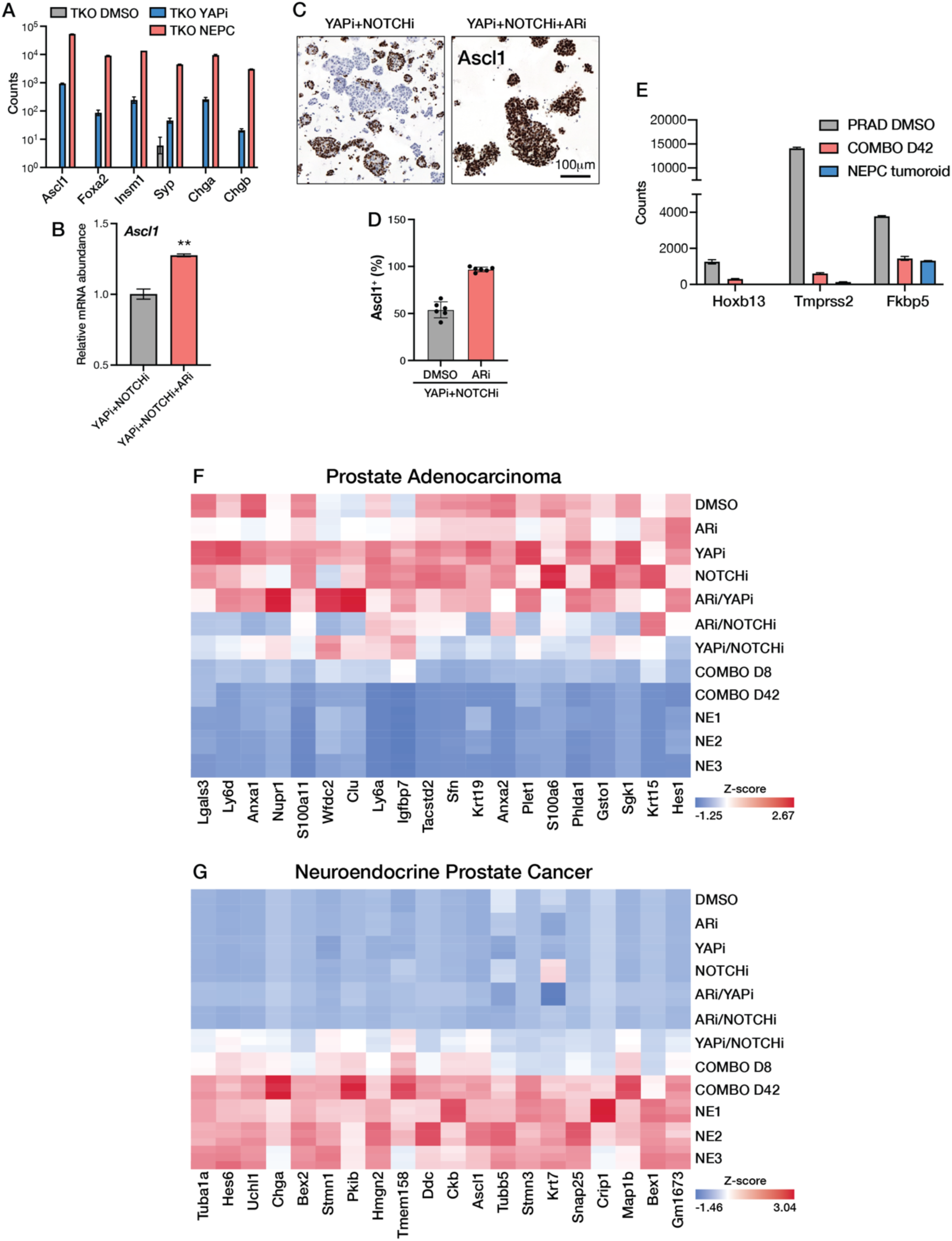
Combined inhibition of YAP/TEAD, NOTCH and AR signaling reprograms PRAD into NEPC. (A) Normalized bulk RNA-seq counts of the indicated NE markers in TKO PRAD organoids treated with DMSO or YAPi (1uM IAG933) for 8 days and in TKO NEPC tumoroids. Error bars, ±SEM; n=3. (B) qRT-PCR analysis showing ARi promotes *Ascl1* induction in the presence of YAPi and NOTCHi following 8 days of treatment. Error bars, ±SEM; n=3; **p<0.01. (C) Representative ASCL1 IHC images of TKO PRAD organoids treated with YAPi+NOTCHi or YAPi+NOTCHi+ARi for 8 days. Scale bar, 100 μm. (D) Quantification of ASCL1-positive cells in TKO PRAD organoids treated with YAPi + NOTCHi in the presence of DMSO or ARi for 8 days. Data are shown as mean ± SEM; n=3. (E) Normalized bulk RNA-seq counts of the indicated PRAD lineage genes in TKO PRAD organoids treated with DMSO or COMBO for 42 days and in TKO NEPC tumoroids. Data are shown as mean ± SEM; n=3. (F) Heatmaps showing expression of prostate adenocarcinoma marker genes across the indicated organoid lines and treatment conditions. (G) Heatmaps showing expression of neuroendocrine prostate cancer marker genes across the same conditions as in (F). The gene sets used in (F) and (G) consist of the top 20 gene from the corresponding gene sets listed in table S2.

**Supplemental Figure 7.**
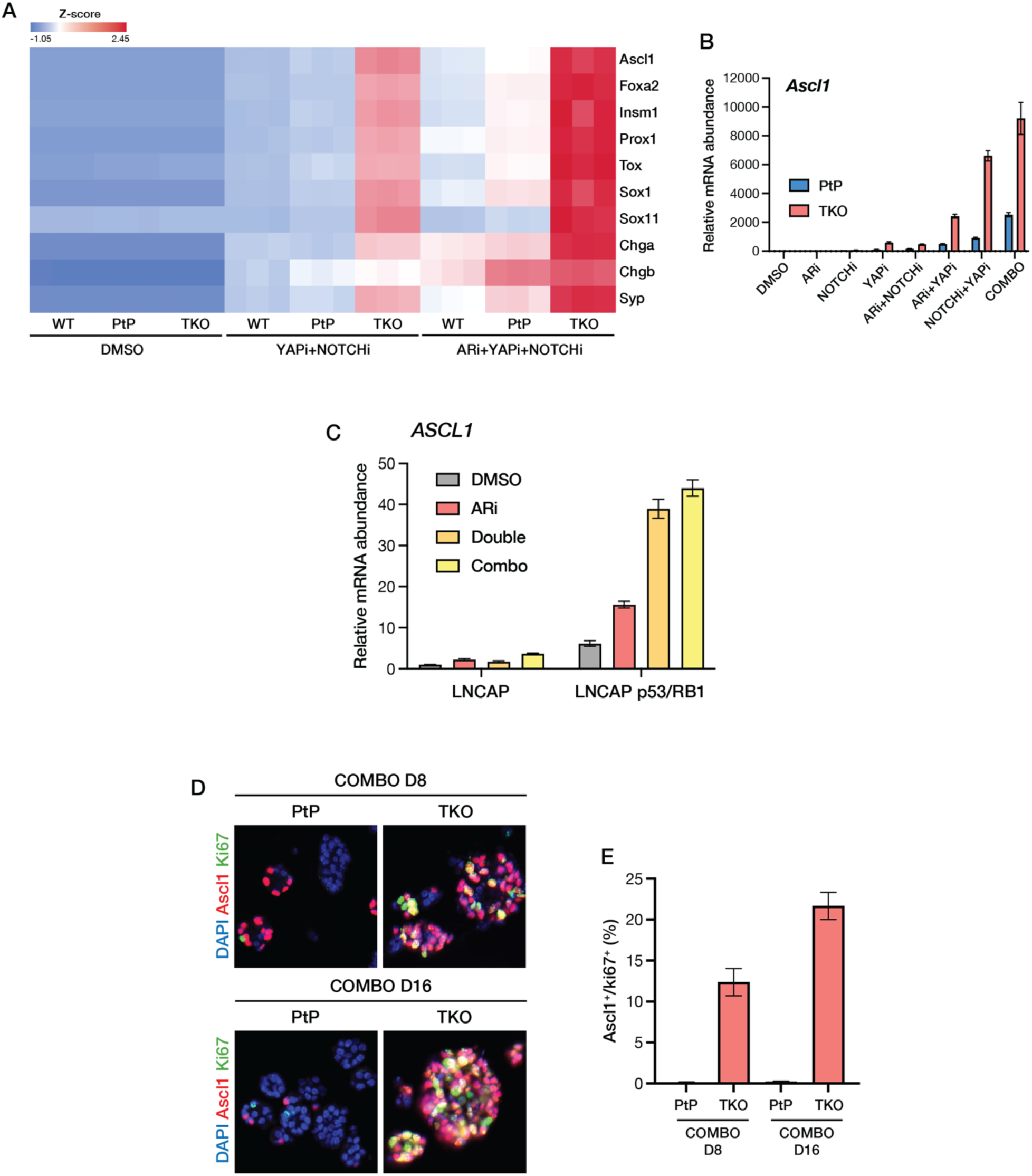
Role of RB1 loss in PRAD-to-NEPC transition. (A) Heatmap showing bulk RNA-seq expression of NE lineage-associated genes in WT, PtP (*Pten^-/-^*, *Trp53*^-/-^) and TKO (*Pten^-/-^*, *Trp53*^-/-^, *Rb1^-/-^*) organoids treated under the indicated conditions. Expression levels are shown as Z-score. (B) qRT-PCR analysis of *Ascl1* expression in bulk PtP and TKO organoids following treatment under the indicated conditions for 8 days. Error bars, ±SEM, n=3. (C) qRT-PCR analysis of *ASCL1* expression in parental LNCaP and LNCaP p53/RB1-deficient cells following treatment under the indicated conditions. Data are shown as mean ± SEM; n=3. (D) Representative immunofluorescence images of ASCL1 (red) and Ki67 (green) in PtP and TKO organoids following COMBO treatment for 8 or 16 days. DAPI (blue) marks nuclei. (E) Quantification of ASCL1/Ki67 double-positive cells in PtP and TKO organoids following COMBO treatment for 8 or 16 days. Data are shown as mean ± SEM.

**Supplemental Figure 8.**
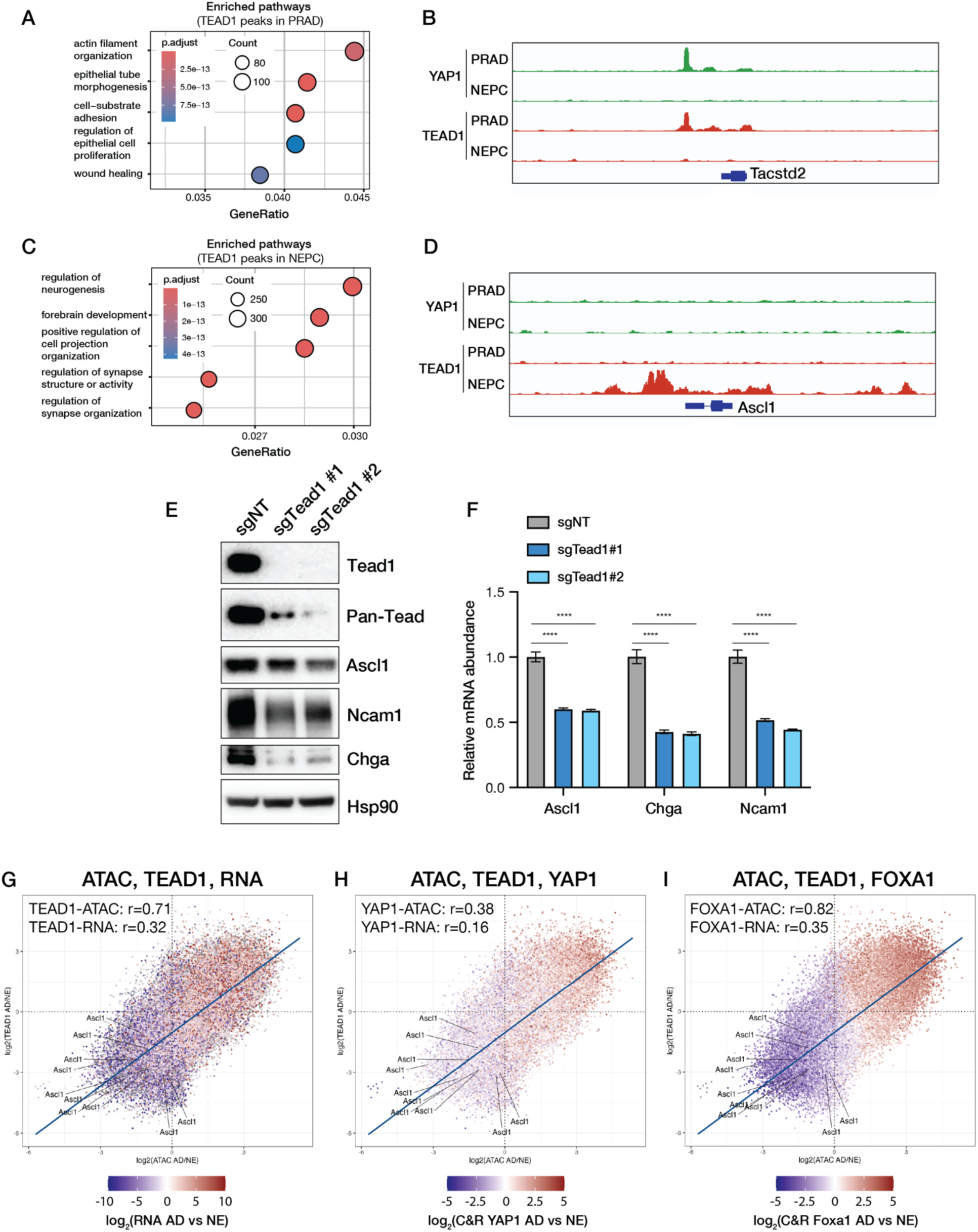
TEAD and FOXA1 are essential for the PRAD to NEPC lineage transition. (A and C) Pathway enrichment analysis of TEAD1 CUT&RUN peaks in PRAD (A) and NEPC (C). Pathways are ranked by gene ratio. (B and D) Genome browser tracks showing CUT&RUN profiles of YAP1 and TEAD1 binding at the adenocarcinoma marker *Tacstd2* (TROP2) and the NE gene *Ascl1* in TKO PRAD and NEPC organoids. Screenshots were generated using Integrative Genomics Viewer (IGV). (E) Western blot analysis demonstrating reduced protein level of NE markers upon Tead1 knockout in TKO NEPC organoids. (F) qRT-PCR showing decreased expression of NEPC markers following Tead1 knockout in TKO NEPC organoids. Error bars represent ±SEM, n=3, ****p<0.0001. (G) Integration of ATAC-seq, TEAD1 CUT&RUN, and bulk RNA-seq profiles comparing PRAD and NEPC states. Regulatory regions are positioned according to differential chromatin accessibility (x-axis) and TEAD1 binding (y-axis) and colored by differential expression of the associated genes. Pearson correlation coefficients between TEAD1 binding and chromatin accessibility or gene expression are indicated. Ascl1-associated regulatory regions are highlighted. (H) Integration of ATAC-seq, TEAD1 CUT&RUN, and YAP1 CUT&RUN profiles comparing PRAD and NEPC states. Regulatory regions are positioned according to differential chromatin accessibility (x-axis) and TEAD1 binding (y-axis) and colored by differential YAP1 binding. Pearson correlation coefficients between YAP1 binding and chromatin accessibility or gene expression are indicated. *Ascl1*-associated regulatory regions are highlighted. (I) Integration of ATAC-seq, TEAD1 CUT&RUN, and FOXA1 CUT&RUN profiles comparing PRAD and NEPC states. Regulatory regions are positioned according to differential chromatin accessibility (x-axis) and TEAD1 binding (y-axis) and colored by differential FOXA1 binding. Pearson correlation coefficients between FOXA1 binding and chromatin accessibility or gene expression are indicated. *Ascl1*-associated regulatory regions are highlighted.

**Supplemental Figure 9.**
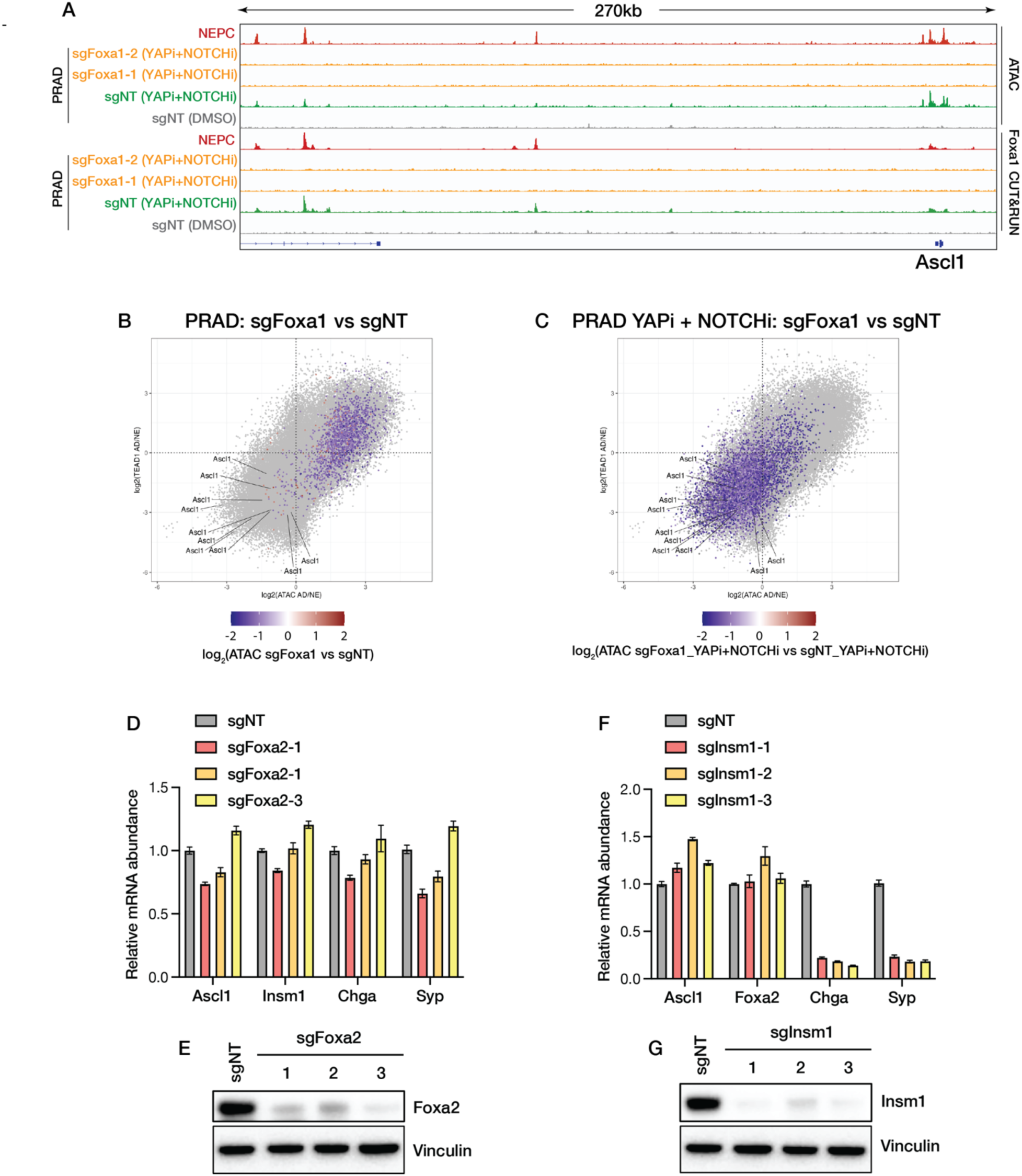
Role of FOXA1, FOXA1 and INSM1 in PRAD-to-NEPC transition. (A) Genome browser tracks showing ATAC-seq and FOXA1 CUT&RUN peaks at the *Ascl1* promoter and enhancer regions across the indicated organoid lines and treatment conditions. Screenshots were generated using IGV. (B) Integration of ATAC-seq and TEAD1 CUT&RUN profiles in PRAD organoids comparing sgFoxa1 with sgNT. Each point represents a regulatory region, positioned according to its differential chromatin accessibility between PRAD and NEPC (x-axis) and differential TEAD1 binding between PRAD and NEPC (y-axis), and colored by the change in chromatin accessibility following *Foxa1* knockout. *Ascl1*-associated regulatory regions are indicated. (C) Integration of ATAC-seq and TEAD1 CUT&RUN profiles in YAPi + NOTCHi-treated PRAD organoids comparing sgFoxa1 with sgNT. Regulatory regions are positioned as in (B) and colored by the change in chromatin accessibility following *Foxa1* knockout under YAPi + NOTCHi treatment. *Ascl1*-associated regulatory regions are indicated. (D) qRT-PCR analysis of NE marker gene expression in sgNT and *Foxa2*-deleted organoids using three independent sgRNAs. Data are shown as mean ± SEM; n=3. (E) Western blot analysis of FOXA2 in sgNT and *Foxa2*-deleted organoids using three independent sgRNAs. Vinculin serves as a loading control. (F) qRT-PCR analysis of NE marker gene expression in sgNT and *Insm1*-deleted organoids using three independent sgRNAs. Data are shown as mean ± SEM; n=3. (G) Western blot analysis of INSM1 in sgNT and *Insm1*-deleted organoids using three independent sgRNAs. Vinculin serves as a loading control.

## Notes

### Competing Interest Statement

Charles L Sawyers serves on the Board of Directors of BeOne and the scientific advisory boards of CellCarta, Column Group, Foghorn, Housey Pharma, Juri, Manas, Nextech, Nilo, PMV Pharma and ORIC.

### Summary of Updates

Figures updated. Include 39 new panels in the figures.

